# Gardenin A improves cognitive and motor function in A53T-α-syn mice

**DOI:** 10.1101/2023.10.27.564401

**Authors:** Wyatt Hack, Noah Gladen-Kolarsky, Swarnali Chatterjee, Qiaoli Liang, Urmila Maitra, Lukasz Ciesla, Nora E Gray

## Abstract

Oxidative stress and neuroinflammation are widespread in the Parkinson’s disease (PD) brain and contribute to the synaptic degradation and dopaminergic cell loss that result in cognitive impairment and motor dysfunction. The polymethoxyflavone Gardenin A (GA) has been shown to activate the NRF2-regulated antioxidant pathway and inhibit the NFkB-dependent pro-inflammatory pathway in a *Drosophila* model of PD. Here, we evaluate the effects of GA on A53T alpha-synuclein overexpressing (A53TSyn) mice.

A53TSyn mice were treated orally for 4 weeks with 0, 25, or 100 mg/kg GA. In the fourth week, mice underwent behavioral testing and tissue was harvested for immunohistochemical analysis of tyrosine hydroxylase (TH) and phosphorylated alpha synuclein (pSyn) expression, and quantification of synaptic, antioxidant and inflammatory gene expression. Results were compared to vehicle-treated C57BL6 mice.

Treatment with 100 mg/kg GA improved associative memory and decreased abnormalities in mobility and gait in A53TSyn mice. GA treatment also reduced cortical and hippocampal levels of pSyn and attenuated the reduction in TH expression in the striatum. Additionally, GA increased cortical expression of NRF2-regulated antioxidant genes and decreased expression of NFkB-dependent pro-inflammatory genes. GA was readily detectable in the brains of treated mice and modulated the lipid profile in the deep gray brain tissue of those animals.

While the beneficial effects of GA on cognitive deficits, motor dysfunction and PD pathology are promising, future studies are needed to further fully elucidate the mechanism of action of GA, optimizing dosing and confirm these effects in other PD models.

**Significance Statement:** The polymethoxyflavone Gardenin A can improve cognitive and motor function and attenuate both increases in phosphorylated alpha synuclein and reductions in tyrosine hydroxylase expression in A53T alpha synuclein overexpressing mice. These effects may be related to activation of the NRF2-regulated antioxidant response and downregulation of NFkB-dependent inflammatory response by Gardenin A in treated animals. The study also showed excellent brain bioavailability of Gardenin A and modifications of the lipid profile, possibly through interactions between Gardenin A with the lipid bilayer, following oral administration. The study confirms neuroprotective activity of Gardenin A previously reported in toxin induced *Drosophila* model of Parkinson’s disease.

## Introduction

Neurodegenerative diseases (ND) such as Parkinson’s disease (PD), along with other non-communicable aging-associated diseases, have become one of the greatest medical challenges of the 21st century. PD is characterized by loss of dopaminergic neurons in the nigrostriatal pathway and an accumulation of Lewy bodies made up of abnormally aggregated alpha synuclein (1). Impaired motor function is the primary clinical manifestation of the disease, although cognitive deficits are also quite prevalent as the disease progresses, affecting as many as 80% of patients who have had the disease 15 years or longer (2). Oxidative stress and uncontrolled neuroinflammation have been implicated as major contributing factors for the development and progression of PD (3) and are thought to promote alpha synuclein accumulation (3-5).

Although PD is the second most diagnosed ND it has limited pharmacological treatment options, providing some relief for motor symptoms while largely ignoring cognitive symptoms (6). Standardized care involves the use of drugs that increase dopaminergic neurotransmission in the midbrain through different mechanisms (6). Levodopa (L-DOPA) is the most prescribed drug for the management of PD because it is converted to dopamine through decarboxylation in the brain (7). Molecules, such as bromocriptine, apomorphine, pergolide and lisuride also modulate dopamine signaling through agonism of the D2 dopaminergic receptor (8). Notably, all drugs named in this paragraph are plant-derived molecules, underscoring how nature can serve as a valuable source of pharmacologically active drug-like molecules (9).

L-DOPA was first identified in broad beans, one of several bean types considered as staples of the Mediterranean diet (10). The Mediterranean diet, rich in vegetables, fruit, and herbs, has been associated with lowering oxidative stress and neuroinflammatory markers in animal studies and human clinical trials (11). In fact, in a small case-control study, adherence to the Mediterranean diet was associated with lower risk of PD, and lower adherence was associated with earlier age of PD onset (12). Many of the beneficial effects of the Mediterranean diet have been linked to a large class of dietary polyphenols: flavonoids (13, 14).

Flavonoids and other dietary phytochemicals have long been disregarded as molecules worth pursuing as potential drugs due to their soft electrophilic character, which makes them inherently reactive molecules (9, 15, 16). High reactivity, often observed in *in vitro* assays, has resulted in these compounds being labelled as pan-assay interference compounds (PAINs) and improbable metabolic panaceas (IMPs) (16, 17). However, the reactivity of these compounds reported using *in vitro* tests does not necessarily translate directly to their biological effects in the context of a living organism (9). It is important, therefore, that biological effects of these phytonutrients are evaluated using whole animal models *in vivo*.

Because of the beneficial link between neuronal health and a diet rich in flavonoids, we attempted to evaluate neuroprotective potential of these molecules first in a toxin-induced *Drosophila* model of PD (15, 18). The screen showed that flavones, a class of flavonoids, lacking a functional group substitution at the C3 position, to be the most neuroprotective (15). Lipophilic polymethoxyflavones were amongst the most biologically active compounds tested in *Drosophila* PD model. Among these molecules, Gardenin A (GA) stood out as one of the most neuroprotective compounds (18).

GA is found in the gum of the medicinal plant, *Gardenia resinifera* Roth (19). It was previously shown to be hepatoprotective and to stimulate neuritogenesis in cell cultures (19, 20). It was also reported to exhibit multiple neuropharmacological effects in mice including sedative, anxiolytic, antidepressant, and anticonvulsant activities (21). Our group’s detailed studies of this molecule in a *Drosophila* model of PD showed that GA exerts pleiotropic effects through the regulation of nuclear factor erythroid 2–related factor 2 (NRF2) as well as the immune deficiency pathway involving NFκB and cellular death responses (18).

The studies presented here build on that previous research to evaluate the effects of GA in a mammalian model of PD. We have employed the A53T alpha synuclein (A53TSyn) transgenic mouse model of alpha synuclein aggregation linked to PD pathogenesis. We found that four weeks of treatment with a high dose GA (100 mg/kg), administered by oral gavage, attenuated motor and cognitive deficits apparent in the A53TSyn mice compared to control group with vehicle alone. GA also increased expression of synaptic and antioxidant genes in the cortex of treated A53TSyn mice and decreased expression of pro-inflammatory genes. Using ultra-performance liquid chromatography (UPLC) with Multiple Reaction Monitoring (MRM) and high-resolution mass spectrometry (HRMS), we also quantified the GA level and characterized major lipid profile modifications in brain tissue from the treated mice.

## Results

### GA improves associative memory in A53TSyn mice

We employed the Conditioned Fear Response (CFR) test to assess contextual memory (Figure 1A). In order to account for potentially confounding effects of mobility impairments, freezing during the test phase is adjusted for baseline freezing levels. We observed that A53TSyn mice had significantly reduced freezing relative to wild type (WT) animals and that treatment with 100mg/kg GA significantly attenuated this deficit (Figure 1B). There was a similar trend towards improved CFR performance in A53TSyn mice with 25mg/kg GA treatment although it did not reach statistical significance.

**Figure 1.**
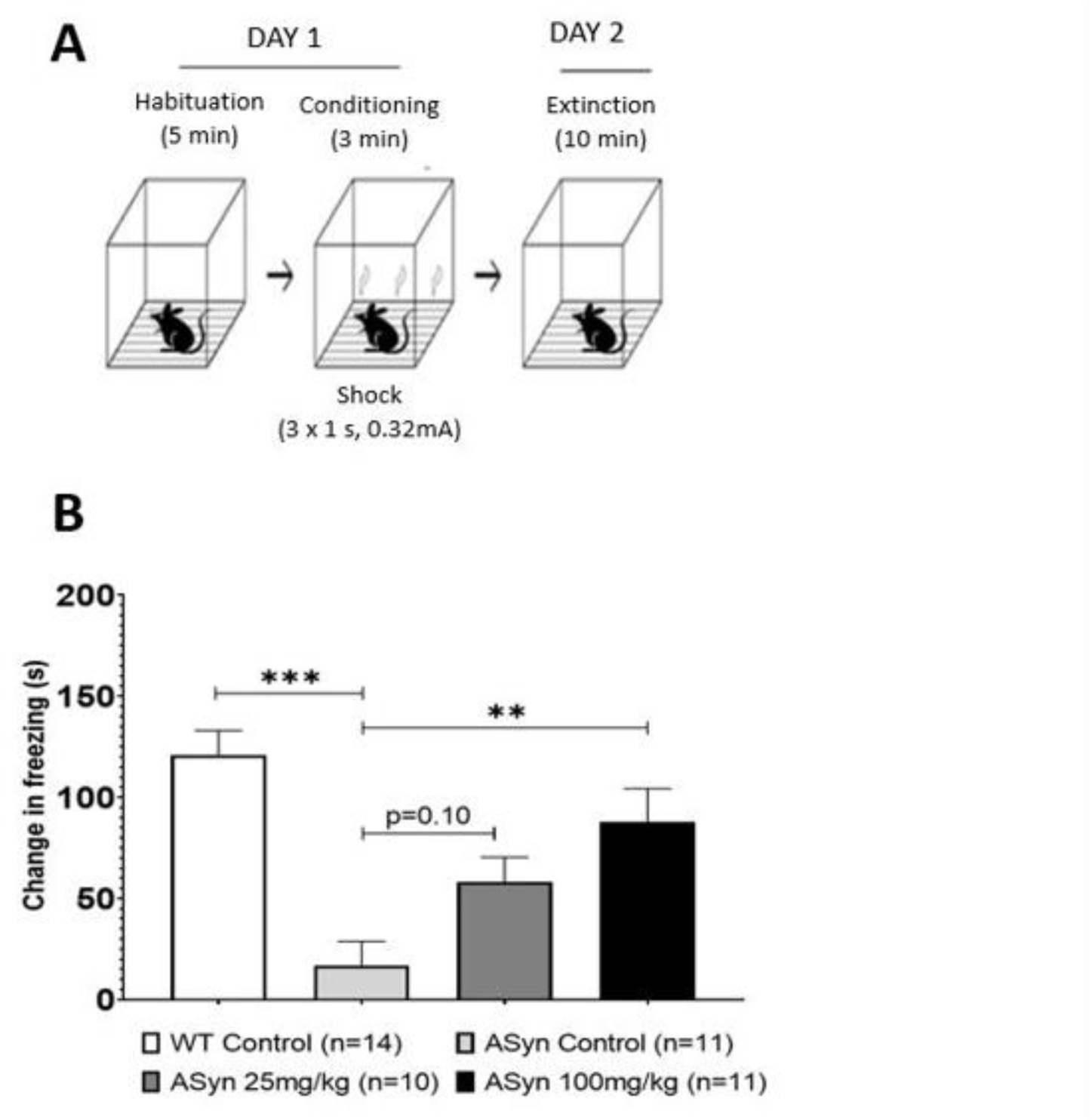
GA attenuated deficits in associative memory in A53TSyn mice. A53TSyn mice spent significantly less time freezing relative to WT animals. When treated with GA at the 100mg/kg dose, A53TSyn mice showed a significant increase in freezing. **p<0.01, ***p<0.001

There was a trend towards decreased synaptophysin expression in the cortex of A53TSyn mice relative to WT animals and 100mg/kg GA significantly increased this expression in the A53TSyn mice (Figure 2). There was a similar but non-significant, pattern in the expression of post-synaptic density protein 95 (PSD95) in the cortex of the mice.

**Figure 2:**
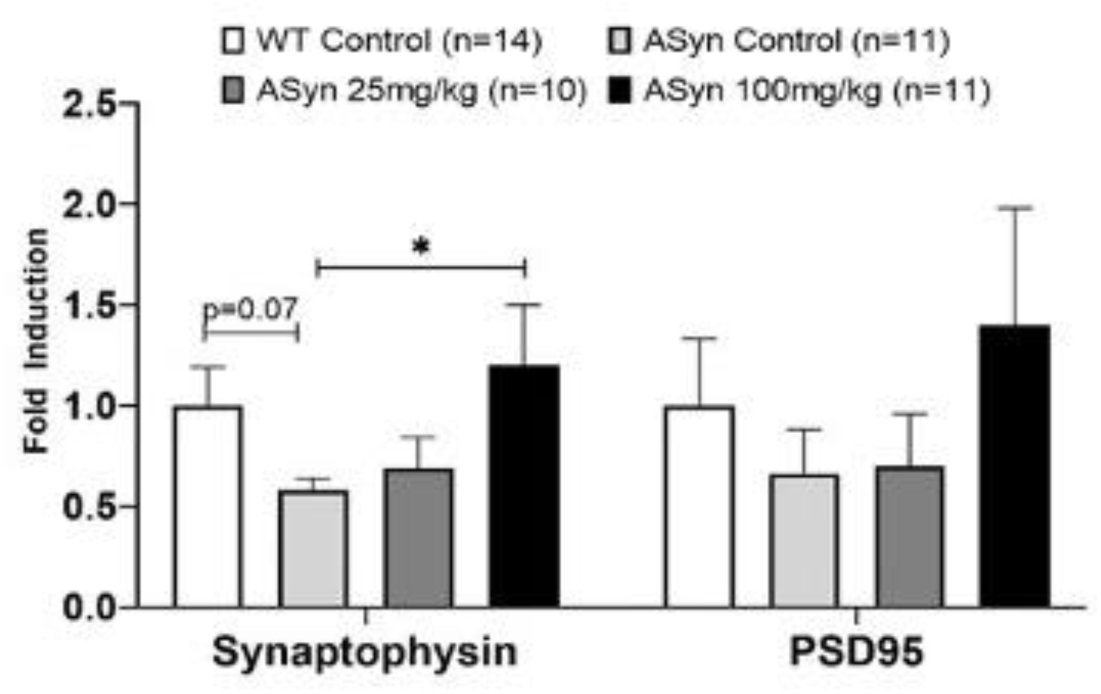
GA increased cortical expression of synaptophysin in A53TSyn mice. A53TSyn mice treated with 100mg/kg GA showed a significant increase in the expression of synaptophysin relative to untreated A53TSyn mice. *p < 0.05

### GA reduces alpha synuclein pathology in the hippocampus and cortex

A53TSyn mice showed robust increases in phosphorylated alpha synuclein (pSyn) relative to WT mice (Figure 3A). 100mg/kg GA significantly reduced expression of pSyn in both the hippocampus (Figure 3B) and cortex (Figure 3C) of A53TSyn mice. Treatment with 25mg/kg GA did not significantly alter pSyn in either brain region (Figure 3B and 3C).

**Figure 3:**
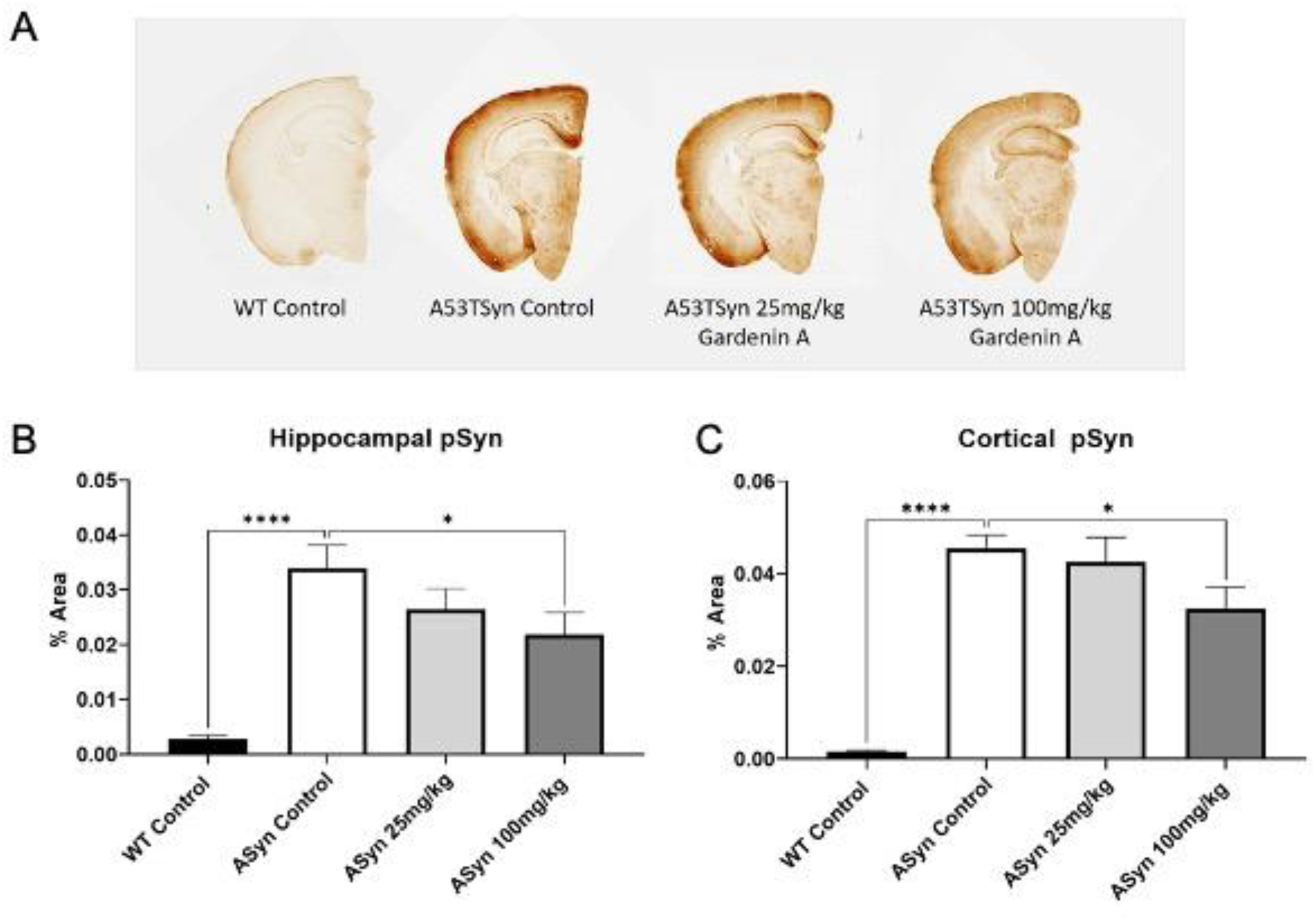
GA decreased pSyn expression in the brain of A53TSyn mice. A) Example images from A53TSyn and WT mice. pSyn expression in the (B) hippocampus and (C) cortex was attenuated in A53TSyn mice treated with 100mg/kg GA. *p<0.01, ****p<0.0001.

### GA improves motor deficits in A53TSyn mice

A53TSyn mice displayed a significant reduction in overall mobility in the open field test relative to WT animals (Figure 4). Treatment with 100mg/kg GA significantly increased mobility in A53TSyn mice to levels that were comparable to the mobility of WT mice (Figure 4). Using the Digigait apparatus to assess temporal and spatial gait differences, we detected alterations in stance width and gait in the A53TSyn mice (Figure 5A and B). Compared to WT animals A53Tsyn mice had increased front paw stance width (Figure 5A). There was a trend towards reduced stance width in A53TSyn mice that were treated with 100mg/kg but it did not reach significance. However, A53TSyn mice treated with 100mg/kg GA no longer showed significant differences in stance width relative to WT mice. There was no effect 25mg/kg GA on front paw stance width in A53TSyn mice nor were any differences were detected between genotypes or treatment groups in the rear paws.

**Figure 4.**
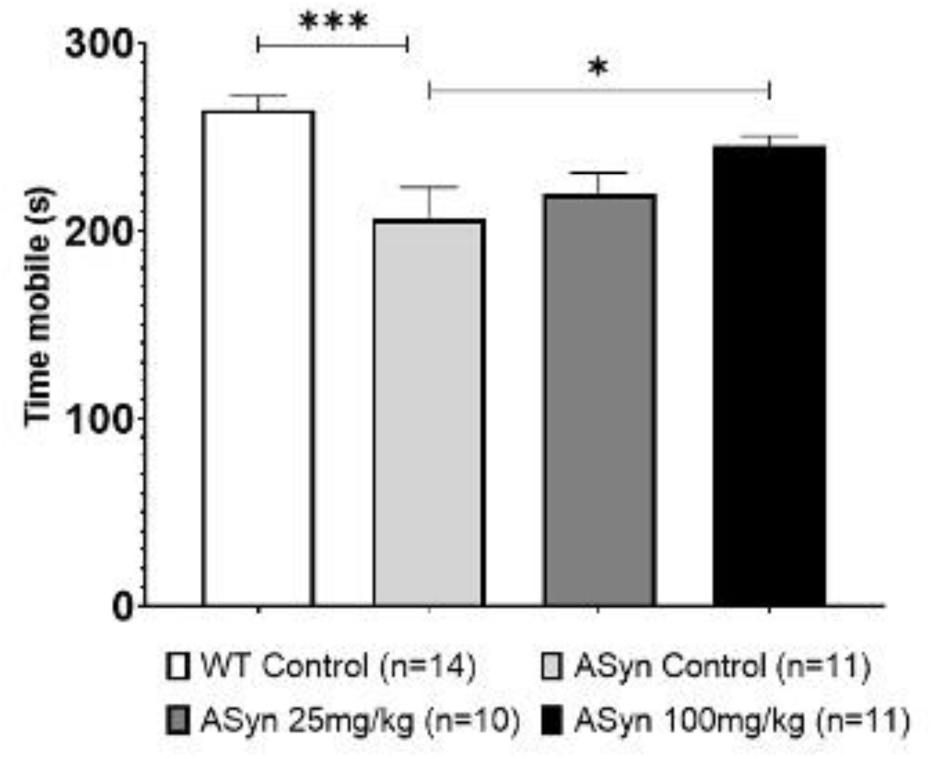
GA attenuated deficits in overall mobility in A53TSyn mice. Treatment with 100mg/kg of GA significantly attenuated deficits in overall mobility compared to untreated A53TSyn mice. *p<0.05, ***p<0.001.

**Figure 5.**
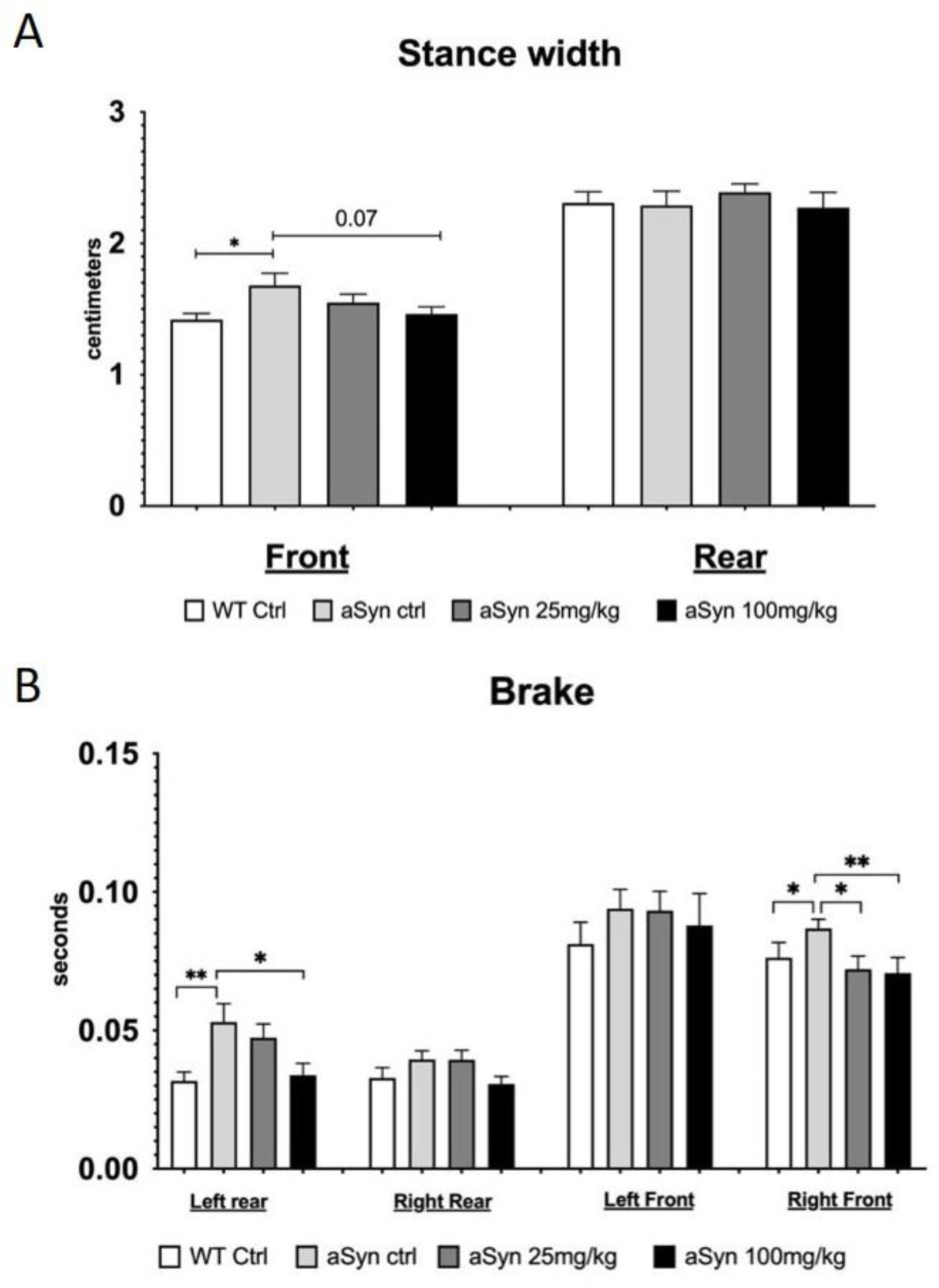
GA attenuated gait abnormalities in A53TSyn mice. A) A53tSyn mice show significantly increased stance width in the front paws compared to WT mice. The front paw stance width of A53TSyn mice treated with 100mg/kg GA was not different from WT mice. B) 100mg/kg GA attenuated increases in brake in the left rear and right front paws in A53TSyn mice. *p<0.05, **p<0.01

Brake was also significantly increased in the A53TSyn mice in the left rear and right front paws (Figure 5B) and this increase was significantly attenuated by 100mg/kg GA treatment. In the right front paw 25mg/kg GA also significantly reduced brake. A similar but non-significant trend towards increased brake in A53TSyn mice and a reduction with the high dose of GA was observed in the right rear and left front paws.

### GA increases tyrosine hydroxylase (TH) expression in the striatum

There was a significant reduction in TH expression in the striatum of A53TSyn mice compared to WT control (Figure 6A). There was a trend towards an attenuation of this reduction in A53TSyn mice treated with 100mg/kg GA (Figure 6B). Although this treatment effect did not reach statistical significance, the levels of TH in the striatum of the 100mg/kg GA treated mice were not lower than WT levels.

**Figure 6.**
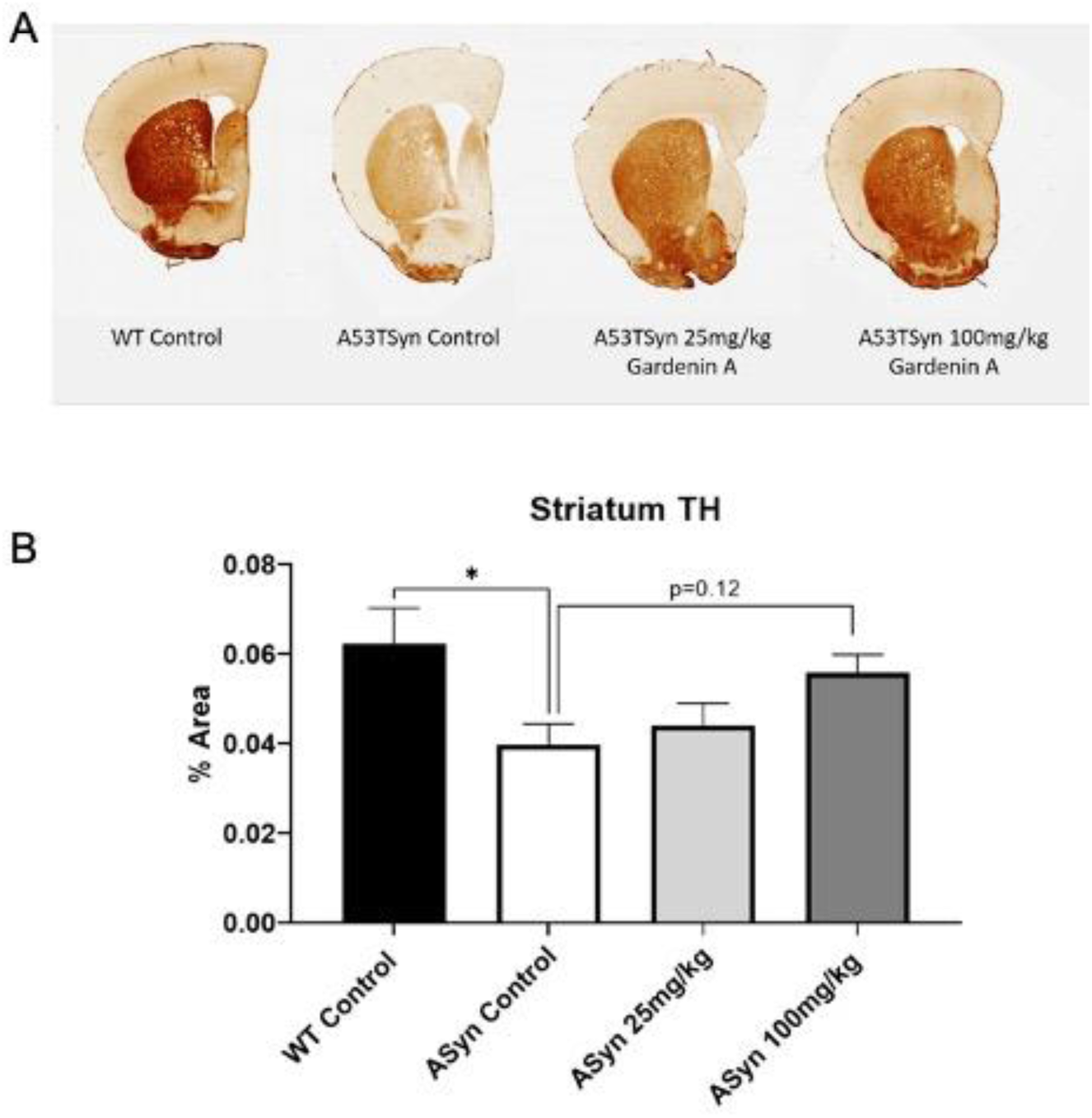
GA restored expression of TH in the striatum of A53TSyn mice. A) Example images from A53TSyn and WT mice B) Control A53TSyn mice showed a significant reduction of TH in the striatum while the expression of TH in A53TSyn mice treated with 100mg/kg GA was not significantly different from WT. *p<,0.05

### GA increases expression of antioxidant genes and decreases expression of pro-inflammatory genes in the cortex of A53TSyn mice

Treatment with 100mg/kg GA increased the expression of NRF2 and its antioxidant target genes in the cortex of A53TSyn mice (Figure 7A). A similar but non-significant trend was observed with 25mg/kg GA treatment.

**Figure 7.**
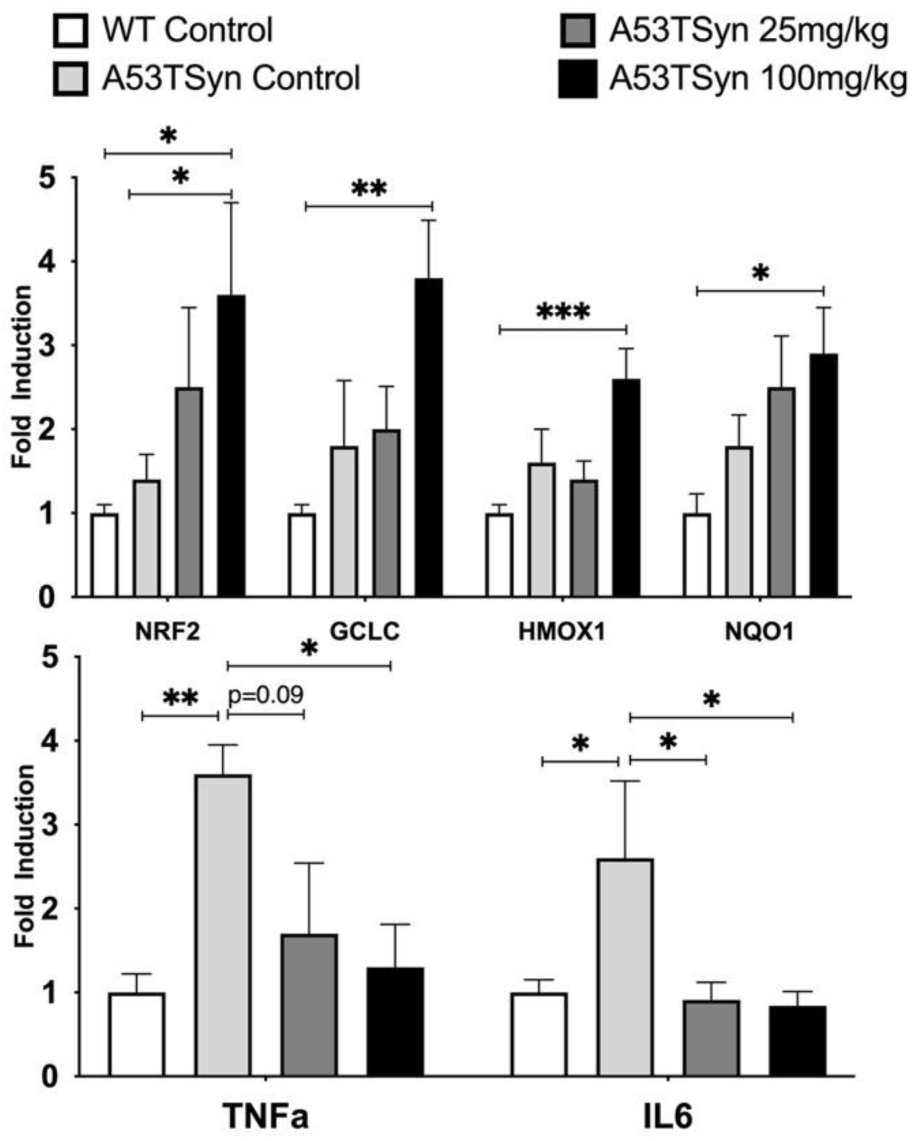
GA treatment modulates expression of antioxidant and inflammatory genes in the cortex of A53TSyn mice. A) Treatment with 100mg/kg of GA significantly increased cortical expression of NRF2 and its target antioxidant genes in A53TSyn mice. B) Treatment with 100mg/kg GA significantly attenuated elevation in expression of pro-inflammatory genes. *p<0.05, **p<0.01, ***p<0.001

Cortical expression of tumor necrosis factor α (TNFα) and interleukin 6 (IL6) was significantly increased in A53TSyn mice. This increase in IL6 was attenuated in A53TSyn mice treated with both 25 mg/kg and 100mg/kg GA (Figure 7B). The increase in TNFα was likewise attenuated with 100mg/kg GA and the 25mg/kg GA dose elicited a similar but non-significant effect on TNFα.

### GA penetrates the blood brain barrier and changes the lipid profile in the deep gray of A53TSyn mice

We assessed GA brain bioavailability using UPLC-MRM. Deep gray tissue was extracted for analysis using the optimized protocol. Nobiletin, a structurally related polymethoxyflavonoid was used at a fixed dose (10 nM) as an internal standard during the extraction procedure. The chromatogram profiles in Figure 8 show the peak of GA and nobiletin at the retention times of 6.94 min; m/z 419.13 and 5.43 ± 0.06 min; m/z 403.14, respectively. We employed the peak integration method to calculate the peak area ratios for GA and nobiletin. We quantified the amount of GA in mice brains based on the standard calibration curve with the peak integration method and linear regression coefficients greater than 0.992. GA was not detected in any of the samples obtained from animals treated with vehicle alone. Nobiletin peaks were present in all chromatograms representing control samples with the recovery of 96.873 ± 2.83% (n = 3), suggesting the optimized extraction method was highly efficient. GA was detected in all brain samples extracted from animals treated with either dose of GA and the data for animals fed with both doses are presented in Figure 8 and Supplementary materials. Interestingly, the concentration of GA in the deep gray tissue reached similar levels for all animals regardless of the dose administered (25 or 100 mg/kg) to the mice (Figure 8 C).

**Figure 8:**
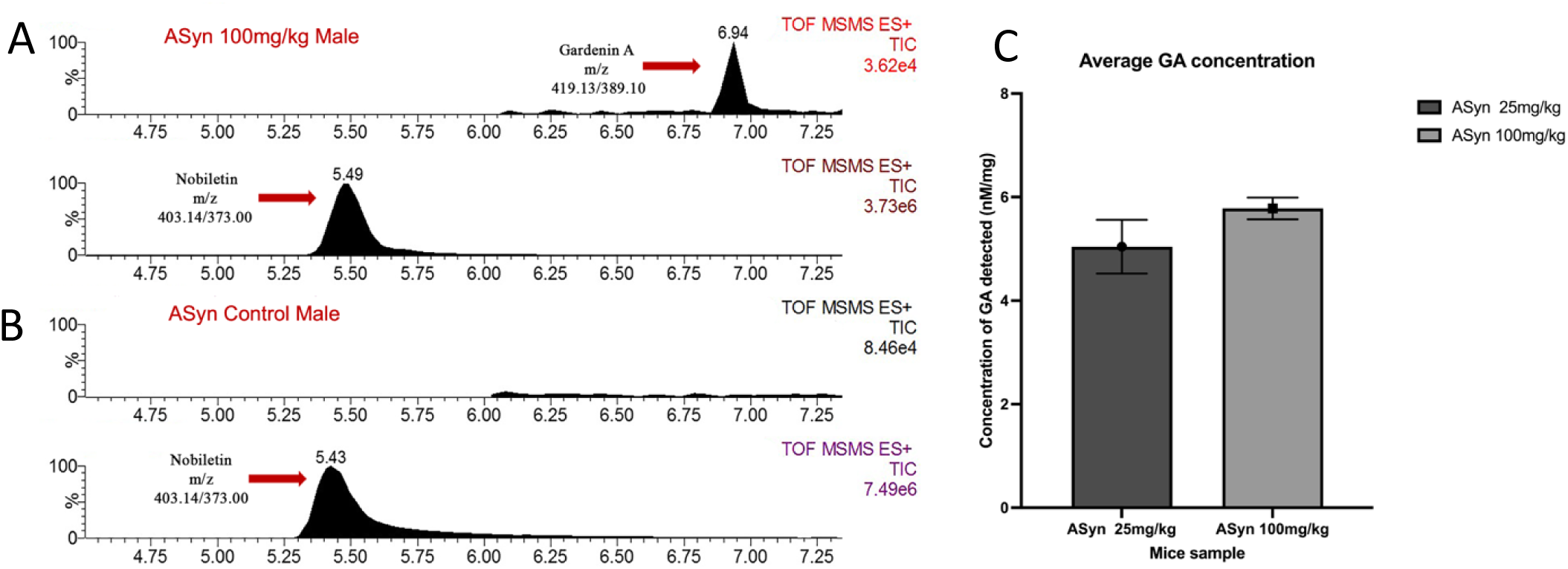
GA bioavailability in A53TSyn mouse brain tissues. (A) Example chromatogram presenting nobiletin and GA peaks in the brain sample obtained from a male mouse treated with 100 mg/kg GA. (B) Example chromatogram presenting the peak of nobiletin and the absence on GA peak in the brain sample obtained from a male mouse treated with a vehicle. (C) Calculated concentration of GA in animals’ brains treated with 25 and 100 mg/kg GA.

Because of this surprising finding, we proceeded to identify possible variations in the GA metabolites and lipids profiles of these samples by searching the HRMS accurate masses (< 5 ppm error) against the previously reported GA metabolite accurate masses and Lipid Maps database (22, 23). Although the GA metabolite search did not generate matches, the lipidomic search revealed the presence of 4 potentially modified lipids which were present only in the male mice brains fed with 100 mg/kg GA. Additionally, we could identify two possible ceramide derivatives, the more saturated one being downregulated and the more unsaturated one being upregulated upon GA administration (Figures 9 and 10 and Table 1), with GA 100 mg/kg treatment showing restoration of the lipid profile close to that of the WT control. Although the HRMS based lipid identification is tentative, the results provide a compelling snapshot demonstrating that GA treatment led to distinct modifications at lipidomic level and suggest the possible interaction of GA or its metabolites with lipids bilayers.

**Figure 9:**
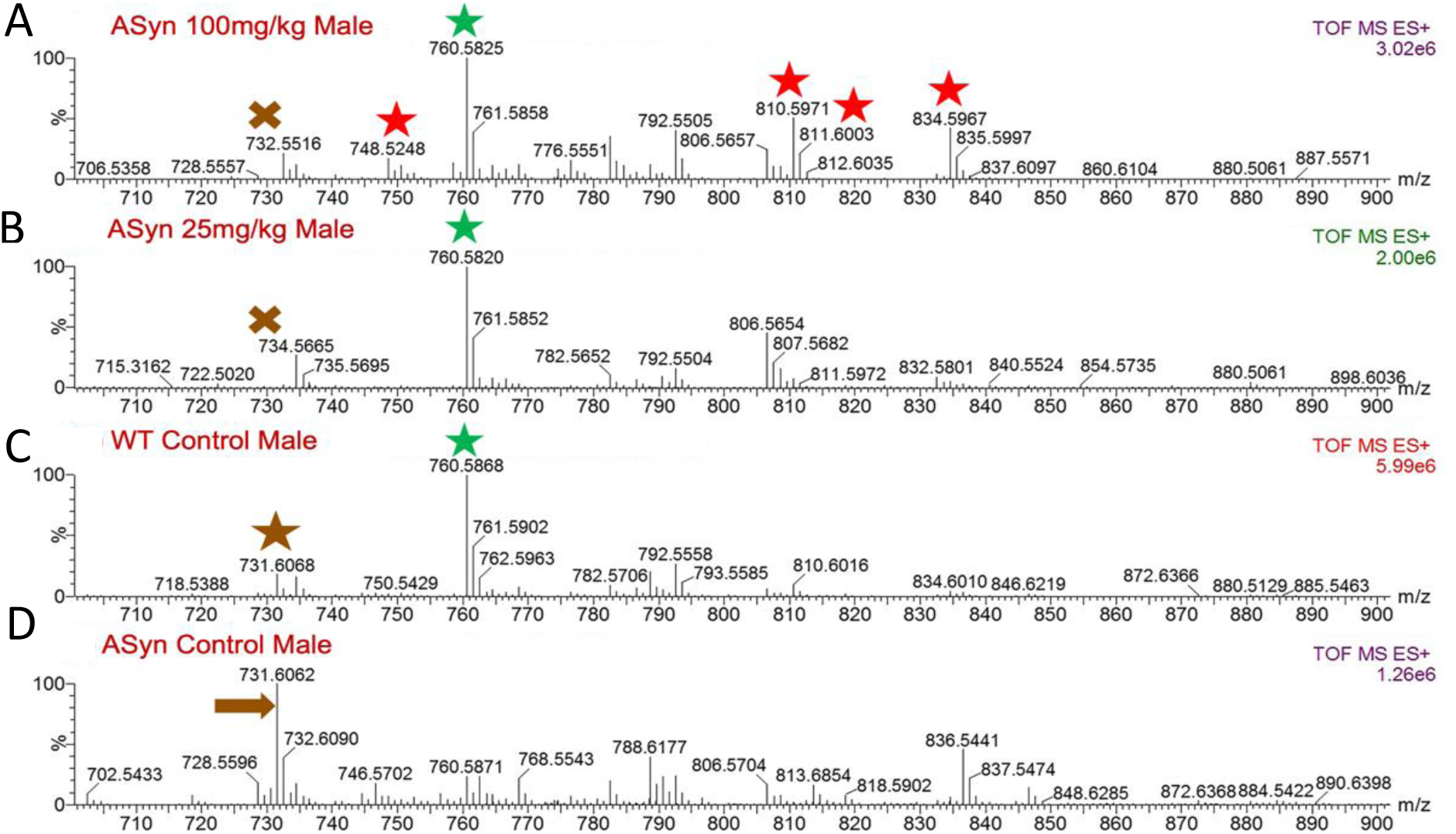
The MS spectra averaged over the HRMS TIC trace 8.0-10.0 min lipid elution region, presenting the detected lipid profile modifications in (A, B) male mouse brains treated with 100 mg/kg and 25 mg/kg GA and (C, D) the *wild type* and *A53TSyn* controls. Lipid ions presented in red were not detected in samples obtained from animals fed with lower dose (25 mg/kg) or in the controls. Ions denoted with green, is a ceramide restored and upregulated in the aSyn mouse brain on oral administration of GA. Ions denoted in brown is another ceramide downregulated in male mice brains after GA administration. Please refer to Table 1 for putative lipid ion assignments.

**Figure 10:**
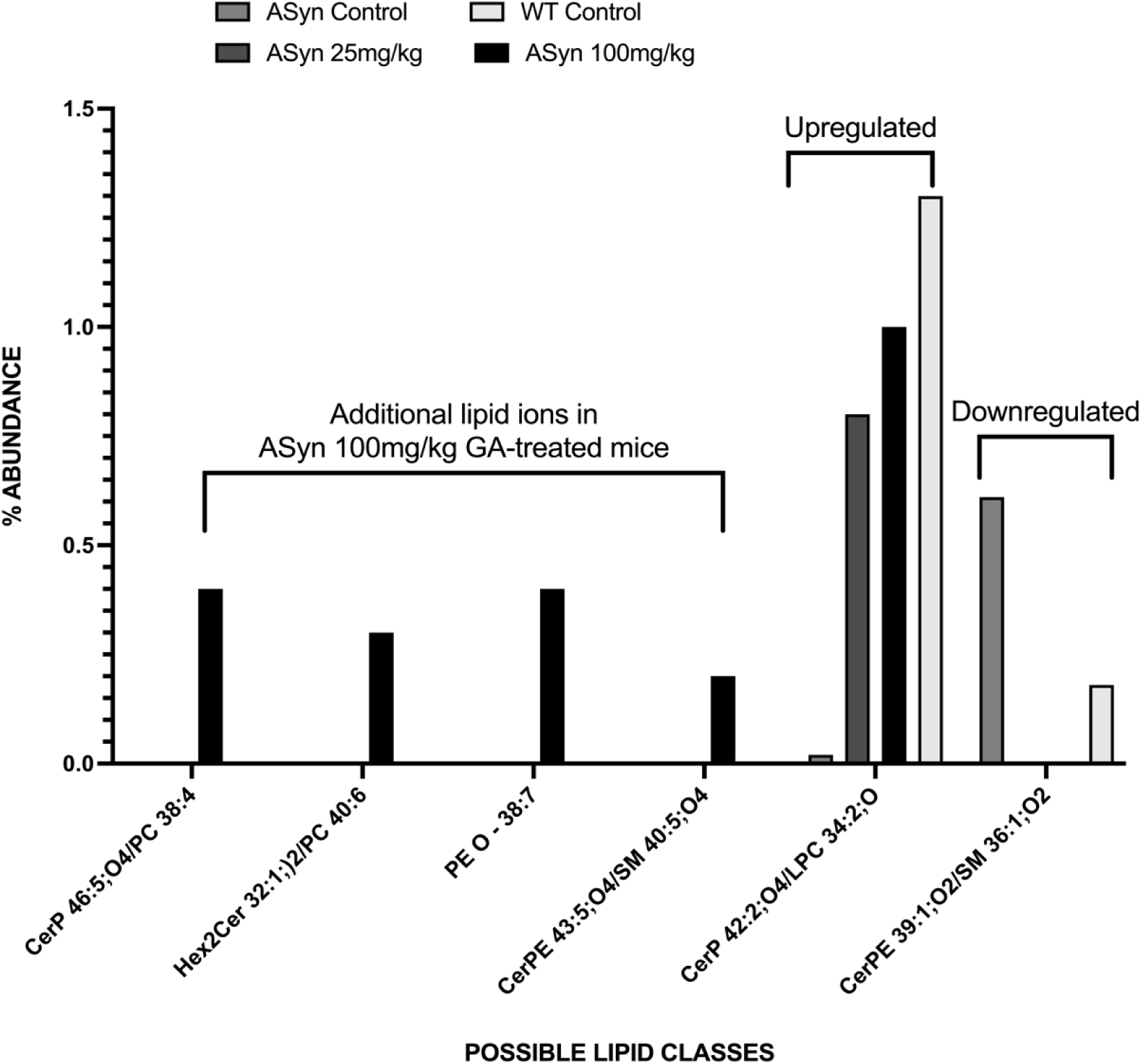
GA modifies lipid profile consisting of ceramides, sphingomyelin, phosphoethanolamine, and/or phosphatidylcholines in the deep gray of A53TSyn mice. Treatment with 100 mg/kg GA significantly modified the lipid profile of the mice cortical brain tissues. Please refer to Table 1 for symbols and putative lipid ion assignments.

**Table 1:**
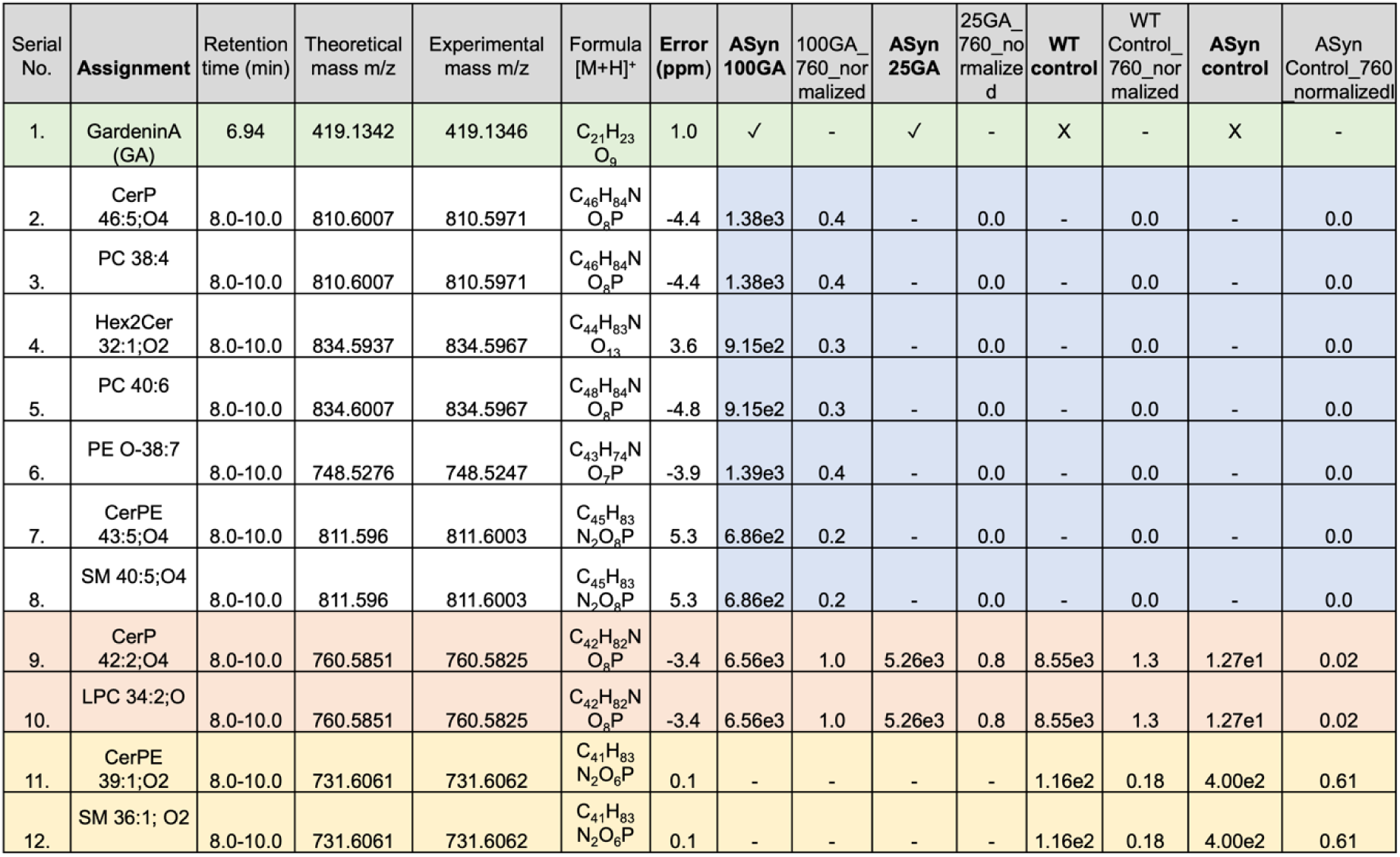
HRMS based putative assignments for GA and lipids detected at retention time 8.0-10.0 mins along with the normalized peak intensities for lipid ions. Assignment (1) confirms the GA presence in the treated mice and absence in the control samples. Assignments (2)-(7) are only present in male mice brains fed with 100mg/kg GA. Assignments (8)-(9) are upregulated and (10)-(11) are downregulated in GA fed male mice brains.

## Discussion

Our data showed that four-weeks of treatment with the polymethoxyflavone GA significantly improved the cognitive and motor impairments in A53TSyn mice. We also found that GA decreased cortical and hippocampal pSyn expression and attenuated deficits in striatal TH expression. Furthermore, GA resulted in increased cortical expression of antioxidant genes and reduced expression of pro-inflammatory genes in treated A53TSyn mice.

Although GA has not been widely studied in mouse models, a recent study published in 2020 reported that doses of GA from 1-25mg/kg in BALB/c mice had potent neuropharmacological effects on anxiety, depression, convulsive response and sleep (21), suggesting central nervous system penetration although it was not directly measured in that study.

While to our knowledge this is the first study of the cognitive effects of GA treatment, the cognitive enhancing effects we observed in this study, along with increased synaptophysin expression, that were elicited by the higher dose of GA are in line with previous reports of the effects of other flavonoids on cognition in human and rodent models. Blueberry extract and isolated compounds from blueberries for instance, have been shown to enhance synaptic plasticity and improve spatial memory in aged rats (24), prevent cognitive impairment induced by mild chronic stress in adult rats (25) and to improve learning and memory in young mice (26). Similar improvements in memory and cognition have been seen in healthy older adults as well as older adults with memory changes (27, 28). The flavonoid epigallocatechin gallate (EGCG), which is found in the tea plant *Camelia sinensis,* has been shown to improve memory and coordination in Alzheimer’s rat models (29, 30), improve mobility and spatial memory in aged mice (31), and exhibit promising effects of reducing stress and fatigue in preliminary human studies (32). Cocoa flavanols have also been shown to have similar improvements in human cognition, increasing verbal fluency, long- and short-term memory, and processing speed in populations of cognitively impaired and cognitively unimpaired older adults (33, 34); additionally, treatment with Acticoa powder, a cocoa extract high in flavonols, showed preventative effects in learning and memory in rats (35, 36), further suggesting the neuroprotective benefits of antioxidants.

In addition to the increases in cognition and synaptic gene expression, we also found that GA treatment decreased the amount of pSyn in the hippocampus and cortex of treated animals. The effects of GA on synuclein pathology have also not been previously reported but other studies of plant extracts and flavonoids have likewise demonstrated effects on alpha synuclein pathology. Extracts of the plant *<u>Scutellaria</u> <u>pinnatifida</u>* have been shown to inhibit alpha synuclein aggregation *in vitro* (37) and tea polyphenols have been found to inhibit alpha synuclein aggregation in the brains of monkeys in an MPTP(1-methyl-4-phenyl-1,2,3,6-tetrahydropyridine) model of PD (38). Additionally, in A53TSyn mice specifically, treatment with grape polyphenols or caffeic acid have also been shown to reduce pSyn pathology (39, 40).

In our study, GA treatment also significantly improved deficits in overall mobility and gait in the A53TSyn mice which was associated with an attenuation of TH loss in the striatum. These effects are similar to what was previously demonstrated by Maitra et al showing that Gardenin A prevents the loss of dopaminergic neurons in a paraquat-induced *Drosophila* model of PD (18). Our results are also in line with the reported effects of other flavonoids in A53TSyn mice. Kuang et al found that a flavonoid-rich extract of the gingko biloba plant improved locomotor activity and rescued TH expression in A53TSyn mice (41). Similarly, when A53TSyn mice were treated with caffeic acid there was an observable reduction in dopaminergic cell loss and increased TH in the striatum (40). Interestingly, the effects we observed on motor function following GA treatment are not consistent with what was reported in the 2020 study of the compounds neuropharmacological effects (21). In that study Alonso-Castro et al. observed no change in the locomotor of male BALB/c mice at 1 or 10mg/kg GA and in fact observed a decrease in rotarod performance at following 25mg/kg GA treatment, although this was only evident at a time point 2h after treatment (21). These divergent results could reflect differences in the sensitivity of Digigait as compared to rotarod or in the response to acute versus chronic treatment with GA or could be due to strain differences in gait and mobility between the healthy BALB/c mice and the A53TSyn model of synucleinopathy.

In this study, we also found some initial evidence of a potential mechanism of action of GA. Cortical expression of the transcription factor NRF2 and its antioxidant target genes was increased in brains of GA treated A53TSyn mice while expression of proinflammatory NFkB target genes was reduced in those same animals relative to vehicle-treated A53TSyn mice. Oxidative stress and neuroinflammation are known to contribute to neuronal loss as well as cognitive and motor deficits in PD (42). The findings from this study are consistent with the previously reported effects of GA in paraquat-induced *Drosophila* model of PD (18). In that study oral GA treatment robustly increased the expression of the *Drosophila* orthologs of NRF2 and decreased NFkB. Our results are also in line with the effects of other flavonoids on antioxidant and anti-inflammatory gene and protein expression. The polyphenol resveratrol has been shown to decrease neuroinflammation and oxidative stress in A53T mice (43). Curcumin has been shown to reduce oxidative stress and be neuroprotective in both rat and mouse models of MPTP-induced PD (44, 45) and to lower blood IL-6 and IL-8 levels in streptozotocin-treated rats (46). Similarly, EGCG treatment was also found to reduce serum expression of IL-6 and TNFa in MPTP-treated mice (47) as well as to protect against hypoxia-induced neuroinflammation and oxidative stress via suppression of NFkB and activation of NRF2 (48).

We found that GA was readily detectable in the brains of treated A53TSyn mice although interestingly the concentration of GA did not significantly vary between animals that received the low dose (25 mg/kg) and those that received the high dose (100mg/kg). This was surprising given the behavioral, immunohistochemistry (IHC) and gene expression results which all showed a greater effect of the high GA dose than the lower dose. Some possible explanations of this phenomenon may stem from previously reported induction of certain intestinal cytochrome P450 enzymes by polymethoxyflavonoids (49). The metabolic fate of GA *in vivo* was previously studied in rat liver microsomes (22). Flavonoids’ extensive *in vivo* metabolism has often been cited as one of several factors that led to their exclusion from serious drug discovery endeavors (16). Others have postulated that *in vivo* flavonoid metabolites, resulting from phase I, phase II and microbial metabolism are biologically active and may be responsible for pharmacological effects of these phytonutrients (50, 51). Previous studies also indicated the effects of flavonoids on the metabolism of lipids in mice (52). These studies showed that flavonoids and their metabolites can attach themselves to the lipid bilayer or interact with membrane receptors, which in turn modify the bilayer lipid profile (52-54). The interaction and penetration of polyphenols with lipid bilayers depend on the structures and concentrations of the polyphenols, the compositions of the lipids, and other factors (55). It is possible that, in our study, the measured GA levels represent only the extractable free from of GA present in the deep gray tissues but does not include GA or its metabolites bound to cellular components including lipids and carbohydrates, especially those in the membranous structures.

We attempted to identify GA metabolites in the deep gray tissue of GA treated mice by analyzing the HRMS total ion chromatograms (TIC) obtained for the analyzed brain samples. Unfortunately, the search did not result in the matched peaks of potential GA metabolites, suggesting that future work including more pooled samples may be needed to improve ion detection. Interestingly, the spectra averaged over the 8.0-10.0 min lipid elution region (Figure 9) contained abundant ion information that showed clear differences between the GA-treated and vehicle-treated mice, demonstrating that GA treatment restored the lipidomic profile close to that observed for the WT vehicle-treated mice. Accurate mass search (< 5 ppm error) with Lipid Maps database resulted in putative assignments of one downregulated (m/z 731.6062 CerPE39:1;O2/SM36:1;O2), one upregulated (m/z 760.5825 CerP 42:2;O4/LPC 34:2;O), and four additional lipid ions (m/z 810.5971 CerP 46:5;O4/PC 38:4, m/z 834.5967 Hex2Cer 32;1;O2/PC 40;6, m/z 748.5247 PE O-38:7, m/z 811.6003 SM 40:5;O4/CerPE 43:5;O4) in the brains of 100 mg/kg GA treated mice. The percentage abundance of the identified lipids was normalized to the highest ion intensity in the GA 100 spectrum (m/z 760.58) for better visual presentation (Figure 10). Ion m/z 760.5825 CerP 42:2;O4/LPC 34:2;O, which was highly suppressed in the ASyn controls, was significantly upregulated in the brains of mice fed with GA at levels close to that present in the *wild type* controls. Ion m/z 731.6062 representing a more saturated ceramide (CerPE39:1;O2/SM36:1;O2) was significantly downregulated in the brains of mice fed with GA, indicating the possible role of GA in modifying the profile of saturated lipids. Note that for some lipid ions (e. g. m/z 760.5825 and m/z 810.5971) the HRMS search led to multiple hits within 5 ppm window and not all were listed. Future in-depth lipidomic investigation is needed to clarify the assignments and quantify the lipids of interest.

These findings are in line with previous studies in mice showing the effects of compounds such as xathohumol on the brain and bile ceramides’ profiles and their metabolites (52). While this data does not prove the involvement of any of these lipids in direct biological effects of GA, they provide possible novel directions for exploring the neuroprotective effects of this compound. Future studies can investigate the role of GA on the brain lipid profiles in both *Drosophila* and mouse models of PD.

This study supports previous findings in a *Drosophila* model of PD and shows similar neuroprotective mechanism in a mammalian model. Our findings further support the usefulness of the approach combining an invertebrate model in a prescreening of large library of phytonutrients followed by evaluation in a mammalian model. The fact that GA improves both motor and cognitive deficits in a PD mouse model makes it particularly attractive to pursue as a therapeutic agent given the dearth of therapies with pleiotropic effects that can modify both symptoms. Future studies will focus on exploring pharmacological effects of GA metabolites, the effects of GA on lipid brain profile, as well as a more in-depth investigation of the mechanism of action of this promising therapeutic agent.

## Materials and Methods

### Animal Rearing and Diet

All experiments were conducted in accordance with the NIH Guidelines for the care and Use of Laboratory Animals and were approved by the institutional Animal Care and Use Committee of the Veteran’s Administration Portland Health Care System (VAPORHCS) (IACUC #:<u>4688-21</u>). Breeding pairs of A53Tsyn mice raised on a C57BL6 background and C57BL6 mice were purchased from Jackson Laboratory (Bar Harbor, ME). Animals were housed in a climate-controlled facility with a 12-h light/12-h dark cycle. Animals were fed 5LOB rodent diet (Lab Diet, MO) and provided with water and diet *ad libitum*. Mice were weaned at 21 days, genotyped at two months of age, and group-housed (3–4 per cage) until 5-6 months of age, when experiments commenced.

### GA Treatment

Five- to six-month-old male and female A53TSyn mice were randomized into vehicle- or GA-treated groups, the latter further divided into equal groups receiving a dose of either 25mg/kg or 100mg/kg GA. A group of vehicle treated C57BL6 mice wildtype (WT) were also included as a control (Table 2).

**Table 2.** Animal numbers in each treatment group (total n=46)

|  | **A53T Syn** |  | **C57BL/6** |  |
| --- | --- | --- | --- | --- |
|  | **Female** | **Male** | **Female** | **Male** |
| Vehicle | 5 | 6 | 7 | 7 |
| 25mg/kg GA | 5 | 5 | 0 | 0 |
| 100 mg/kg GA | 5 | 6 | 0 | 0 |

GA (Sigma, St Louis, MO and ABCR, Karlsruhe, Germany) was suspended in 0.5% (w/v) carboxymethyl cellulose (CMC)-physiological saline (vehicle) and given orally 3 times per week by gavage for a total of 4 weeks (Figure 11).

**Figure 11.**
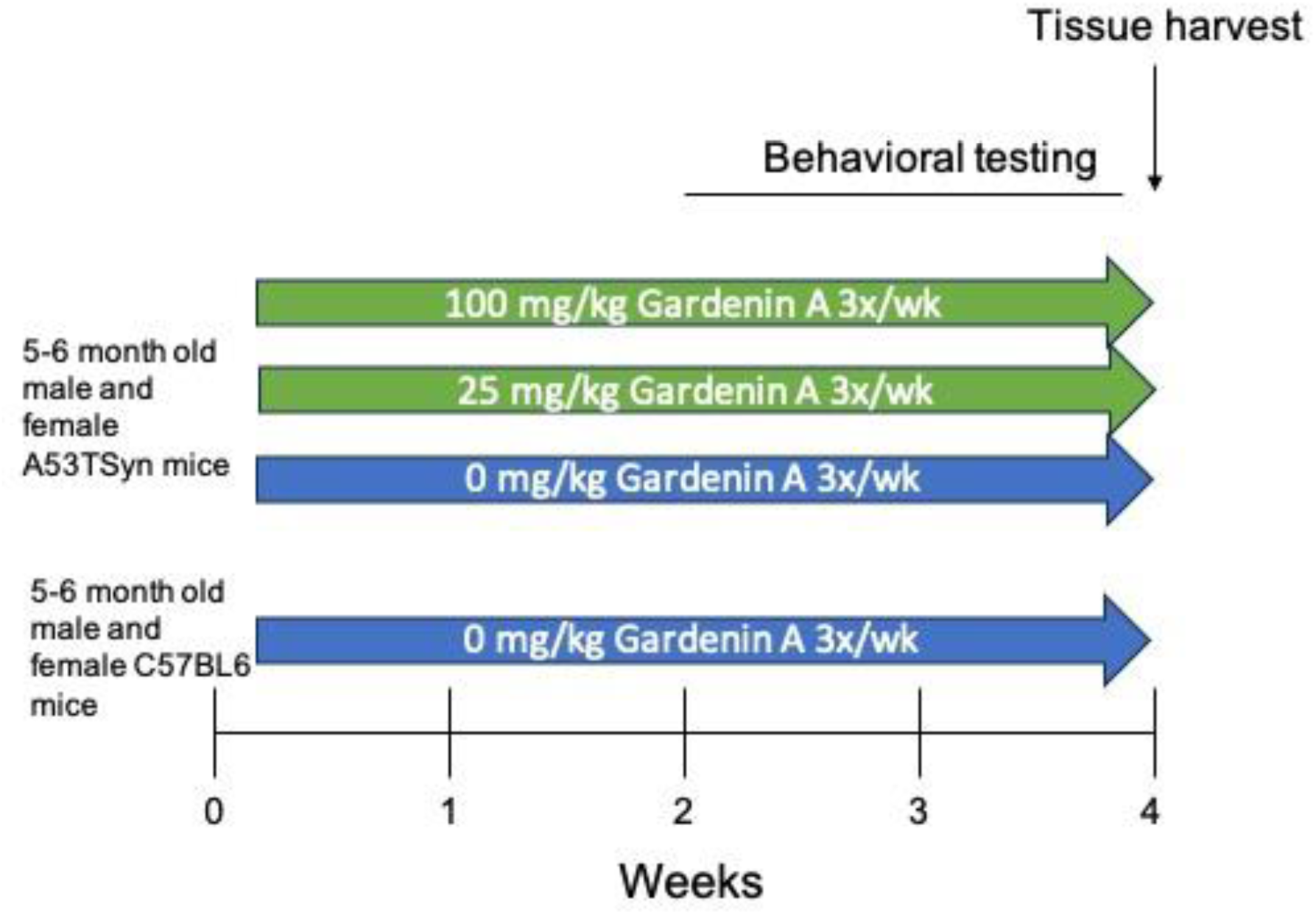
Timeline of experiments.

### Behavioral Analysis

#### Conditioned Fear Response

Conditioned Fear Response (CFR) was conducted over two days to evaluate associative memory. On day 1 mice were allowed a habituation period of 5 minutes as a measure of baseline freezing, followed by three mild shocks (1 s, 0.32 mA) administered to their feet at a random time within a three minute period. The shock stimulus was performed and regulated by Fusion v6.0 for SuperFlex Edition Software (Omnitech Electronics, Inc, Columbus, OH, USA).

After 24 hours, mice were returned to the apparatus for a total of 10 minutes to assess their exhibition of a contextual fear response. No shock is administered during this contextual fear testing stage. Mice were recorded using AnyMaze(Any-Maze Software version 6, Stoelting Co., Wood Dale, IL, USA) and tested on their amount of time freezing, defined as a period of inactivity a minimum of 1000 milliseconds in length. The data is represented as a change in freezing (the difference in freezing on day 2 as compared to day 1) so that differences in overall mobility can be accounted for without confounding the CFR data.

#### Overall mobility

The open field test was conducted over two days in order to test mice on their overall mobility. Each mouse was placed in a square arena (38 × 38 × 64 cm high, constructed of white acrylonitrile butadiene styrene) for 10 min each day and allowed to freely explore the area for environmental acclimatization. The arena was wiped down with chlorhexidine (Novalsan) in between each mouse trial to eliminate scent cues and clean and disinfect the area. Time mice spent mobile was recorded using video tracking software(Any-Maze Software version 6, Stoelting Co., Wood Dale, IL, USA).

#### DigiGait

The DigiGait apparatus (Mouse Specifics, Quincy, MA), was used to assess subtle differences in gait as previously by Goldberg et al (56). Briefly, the DigiGait uses ventral plane videography to capture the gait of each mouse through a transparent, motor-driven treadmill belt. Digital images of the paws of each mouse were taken at 150 frames/sec as mice walk at a speed of 24 cm/sec on the treadmill at an incline of 15 degrees. Each mouse was tested individually on the treadmill, bounded by an acrylic compartment (5 cm in width, 25 cm in length). Temporal and spatial measurements are indicated by the area of the paw relative to the treadmill belt at each frame. Each mouse was recorded for about 3-6 seconds and then the videos were analyzed using the DigiGait software to determine any deficits in several measures of gait and mobility.

### Euthanization and Tissue Collection

After behavioral analysis, animals were anesthetized with inhalable <u>isoflurane</u> and exsanguinated via cardiac puncture. Brain samples and plasma were collected. The right hemisphere was immersion-fixed in 4% <u>paraformaldehyde</u> (Fisher Scientific, MA) in <u>phosphate buffered</u> <u>saline</u> for immunohistochemical analysis and the left hemisphere was sub-dissected and frozen for gene expression analysis by quantitative PCR and GA quantification by UPLC-MRM on a Waters Xevo_G2 XS quadrupole time-of-flight (QTof) instrument.

### Immunohistochemistry

Right hemispheres were incubated in 4% PFA for 24 hours at room temperature. Following PFA incubation, right hemispheres were transferred to 1x PBS and incubated for a further 24 hours, then subsequently incubated in 15% and 30% sucrose solutions before being frozen at -80 C for sectioning. Frozen coronal sections (40µM) were cut on a freezing microtome and stored in a sectioning solution (15% glycerol, 10% TBS, diluted in diH2O) before subsequent processing.

Similar depth sections were placed in a quenching solution (30% methanol, 10% hydrogen peroxide, 10% TBS) for endogenous catalase activity and then blocked (8g bovine serum albumin, 10% horse serum, 2% triton x, 10% TBS, diluted in diH2O). Sections were then incubated with primary antibodies to visualize phosphoryated α-synuclein (pSyn; S129, Abcam) or tyrosine hydroxlyase (TH; Thermo). Following incubation with secondary antibody, Avidin/Biotin Complex) kit (Vector laboratories, Burlingame, CA) was used to visualize staining. DAB (Sigma Fast 3,3 Diaminobenzidine Tablet Set, D-4418 (Sigma-Aldrich Corp, St. Louis, MO, USA)) staining was used as a counterstain.

Sections were then mounted on UltraSlips coverglass slides (GC2450-ACS) and scanned using PrimeHisto XE (Pacific Image Electronics, Torrance CA, USA). ImageJ software (ImageJ, Wayne Rasband, NIH, USA) was used to quantify images. Images were converted to 8-bit grayscale, and the polygon tool was used to encompass the hippocampus or cortex and the total area of each brain area was recorded. The threshold was adjusted to remove ubiquitous background staining, highlighting only areas of intense staining. Staining was quantified in three coronal sections representing regions of anterior, middle and posterior hippocampus and cortex from each right hemisphere sample. Hippocampal and cortical areas were traced using a computerized stage and stereo investigator software. The extent of pSYN or TH staining was expressed as a percentage of the cortical, hippocampal, or striatal area (cm2) that was occupied by detectable immunoreactive staining. Mean values for each parameter were calculated using at least three sections per animal.

### RNA Isolation and Quantitative PCR

Quantitative PCR (qPCR) was performed as described before by Quinn et al(57); briefly, cortical tissue was homogenized in Tri-Reagent (Sigma-Aldrich, MO) and RNA extracted as per the manufacturer’s protocol. Relative gene expression was determined by qPCR using Taqman Gene Expression Master Mix (Thermo Fisher, MA) and commercially available TaqMan primers (Thermo Fisher, MA) on an Applied Biosystems StepOnePlus Real-Time PCR System (Thermo Fisher, MA). All primers were normalized to glyceraldehyde-3phosphate dehydrogenase (GAPDH; Hs02758991_g1). Other primers used were specific for PSD95 (Mm00492193_m1), Synaptophysin (Mm00436850_m1), TNFα (Mm00443258_m1), IL-6 (Mm00446190_m1), NRF2 (Mm00477784_m1), glutamatecysteine ligase catalytic subunit (GCLC -Mm00802655_m1), heme oxygenase 1 (HMOX1 - Mm00516005_m1), and NQO1(Mm01253561_m1). Relative expression was determined using the delta-delta Ct method.

### Quantification of GA in brain tissue

#### Chemicals

Nobiletin (purity ≥ 97%) and GA (purity ≥ 90%) were purchased from Sigma-Aldrich. The identity, purity and stability of the analytical standards was confirmed using an in-house HPLC-MSD system.

#### Biological sample preparation

Approximately 50 mg of deep gray tissue from each mouse was transferred into a conical glass tissue grinder. The tissues were manually ground with a clean pestle in appropriate volume (300 µL per sample) of 60% aqueous acetonitrile (VWR). Internal standard nobiletin (10 nM) was then added at a volume of 0.3 μL per brain sample during the homogenization. The tissue homogenate was centrifuged at 10500 rpm for 15 mins, the supernatant was separated out and filtered through a sterile 0.2 μm cellulose acetate syringe filter.

Prior to mass spectrometric analysis, the filtered solution was further purified and concentrated using Pierce C18 spin columns (Thermo Fisher Scientific) in accordance with the instructions provided by the manufacturer. The clean mixture was subjected to reverse phase UPLC-MRM analysis.

#### UPLC-MRM analysis

The purified and concentrated brain homogenates were subjected to UPLC-MRM analysis using a Waters Xevo G2-XS QTof mass spectrometer connected to an ACUITY I-Class UPLC system. An ACUITY UPLC BEH C18 column (2.1 x 50 mm, 1.7 μm) was used with an ACUITY UPLC BEH C18 Van Guard Pre-column (2.1 x 5 mm, 1.7 μm). The column was eluted at flow rate of 0.3 mL/min with mobile phases A (water with 0.1% formic acid) and B (acetonitrile with 0.1% formic acid) gradient: 0.0 min 80% A 20% B, 5.0 min 60% A 40% B, 10.0-15.0 min 0% A 100% B, 16.0-19.0 min 80% A and 20% B. The injection volume for each sample was 5 μL. MRM analysis was performed in resolution positive ion mode on m/z 403.1→373.0 for nobiletin during 0.0-8.0 min and on 419.1→389.1 for GA during 6.0-19.0 min (collision energy ramp 20-35 eV, target ions enhanced), and simultaneously a HRMS full scan spectrum was recorded (by selecting the RADAR option in MRM method setup), with lockSpray correction on to ensure high accuracy mass measurement. The ESI source conditions included: Capillary voltage 1.23 kV, Sampling cone 40 V, Source offset 80, Source temperature 100 °C, Desolvation temperature 280 °C, Cone gas 15 L/h, Desolvation gas 400 L/h, LockSpray capillary voltage 0.08 kV, Lock mass at 556.2771 in positive ionization mode (leucine enkephalin, peak resolution 28,000). To quantify GA concentrations in the mice brain homogenates, a calibration curve was constructed by plotting the integrated MRM chromatogram peak area ratios (no peak smoothing) of GA calibration standards (1, 10, 25, 50, 75 and 100 nM) and internal standard nobiletin (10nM) versus their concentration ratios. To calculate % recovery, an external calibration curve of Nobiletin was constructed by plotting the integrated chromatogram peak areas (no peak smoothing) versus Nobiletin concentrations (50, 100, 250, 500, 750 and 1000 nM). A blank of ultrapure distilled water was introduced in between each sample to ensure that there was no carryover of the sample in the autosampler. The final concentration of GA in the individual brain samples were estimated based on the total quantities per mg ± SD of the samples for various feeding conditions. After MRM chromatogram quantification of GA from the brain samples, the HRMS full scan TIC averaged spectra at 8.0-10.0 min were plotted for the GA-treated mice samples and controls. Up or down regulated lipid ions were identified and putatively assigned by matching their experimental masses to the theoretical masses in “Lipid Maps” database within 5 ppm error. The ion intensity of the identified lipids was normalized to the highest ion intensity in the GA 100 spectrum (m/z 760.58) for better visual presentation. The integration of a HRMS full scan within an MRM method allowed to both quantify the bioavailability of GA as well as scan for modifications to the lipid profile.

At least three independent biological replicates and two technical replicates per sample were considered for data analysis. The calibration curve was prepared using three replicates per standard dilution.

#### Statistics

All graphs show error bars that reflect standard error of the mean. Statistical significance was calculated using ANOVAs with Sidak pairwise post-hoc testing. Effects were considered significant at *p* ≤ 0.05. Statistically significant differences are indicated in each figure via the following convention: *p < 0.05, \*\**p* < 0.01, \*\*\**p* < 0.001, \*\*\*\**p* < 0.0001. Analyses were performed using GraphPad Prism 8 software (GraphPad Software, Inc., La Jolla, CA).

## Acknowledgments

This study was supported, in part, by the Medical Research Fund through a New Investigator grant (Gray). This work was also supported by the National Center for Complimentary and Integrative Medicine of the National Institutes of Health under awards number 1R41AT011716–01 (Ciesla) and R03AT011871-01 (Ciesla & Maitra). The content is solely the responsibility of the authors and does not necessarily represent the official views of the National Institutes of Health.

## References

1. Jankovic J, Tan EK. Parkinson’s disease: etiopathogenesis and treatment. J Neurol Neurosurg Psychiatry. 2020;91(8):795-808. Epub 2020/06/25. doi: 10.1136/jnnp-2019-322338. PubMed PMID: 32576618.

2. Biundo R, Weise L, Antonini A. Cognitive decline in Parkinson’s disease: the complex picture. NPJ Parkinson’s Disease. 2016;2:16018.

3. Taylor JM, Main BS, Crack PJ. Neuroinflammation and oxidative stress: co-conspirators in the pathology of Parkinson’s disease. Neurochem Int. 2013;62(5):803–19. Epub 2013/01/08. doi: 10.1016/j.neuint.2012.12.016. PubMed PMID: 23291248.

4. Toulorge D, Schapira AH, Hajj R. Molecular changes in the postmortem parkinsonian brain. J Neurochem. 2016;139 Suppl 1:27–58. Epub 2016/07/07. doi: 10.1111/jnc.13696. PubMed PMID: 27381749.

5. Puspita L, Chung SY, Shim JW. Oxidative stress and cellular pathologies in Parkinson’s disease. Mol Brain. 2017;10(1):53. Epub 2017/12/01. doi: 10.1186/s13041-017-0340-9. PubMed PMID: 29183391; PMCID: PMC5706368.

6. Connolly BS, Lang AE. Pharmacological treatment of Parkinson disease: a review. JAMA. 2014;311(16):1670–83. Epub 2014/04/24. doi: 10.1001/jama.2014.3654. PubMed PMID: 24756517.

7. Pereira JB, Kumar A, Hall S, Palmqvist S, Stomrud E, Bali D, Parchi P, Mattsson-Carlgren N, Janelidze S, Hansson O. DOPA decarboxylase is an emerging biomarker for Parkinsonian disorders including preclinical Lewy body disease. Nat Aging. 2023;3(10):1201–9. Epub 2023/09/19. doi: 10.1038/s43587-023-00478-y. PubMed PMID: 37723208; PMCID: PMC10570139 Radiopharmaceuticals, Biogen, Eli Lilly, Eisai, Fujirebio, GE Healthcare, Pfizer and Roche. In the past 2 years, he has received consultancy and speaker fees from AC Immune, Amylyx, Alzpath, BioArctic, Biogen, Cerveau, Eisai, Eli Lilly, Fujirebio, Genentech, Merck, Novartis, Novo Nordisk, Roche, Sanofi and Siemens. S.P. has acquired research support (for the institution) from ki elements/ADDF. In the past 2 years, he has received consultancy and speaker fees from Bioartic, Biogen, Cytox, Eli Lilly, Geras Solutions and Roche. The other authors declare no competing interests.

8. De Keyser J, De Backer JP, Wilczak N, Herroelen L. Dopamine agonists used in the treatment of Parkinson’s disease and their selectivity for the D1, D2, and D3 dopamine receptors in human striatum. Prog Neuropsychopharmacol Biol Psychiatry. 1995;19(7):1147-54. Epub 1995/11/01. doi: 10.1016/0278-5846(95)00232-4. PubMed PMID: 8787038.

9. Maitra U, Stephen C, Ciesla LM. Drug discovery from natural products - Old problems and novel solutions for the treatment of neurodegenerative diseases. J Pharm Biomed Anal. 2022;210:114553. Epub 2021/12/31. doi: 10.1016/j.jpba.2021.114553. PubMed PMID: 34968995; PMCID: PMC8792363.

10. Hornykiewicz O. L-DOPA: from a biologically inactive amino acid to a successful therapeutic agent. Amino Acids. 2002;23(1-3):65–70. Epub 2002/10/10. doi: 10.1007/s00726-001-0111-9. PubMed PMID: 12373520.

11. Aleksandrova K, Koelman L, Rodrigues CE. Dietary patterns and biomarkers of oxidative stress and inflammation: A systematic review of observational and intervention studies. Redox Biol. 2021;42:101869. Epub 2021/02/06. doi: 10.1016/j.redox.2021.101869. PubMed PMID: 33541846; PMCID: PMC8113044.

12. Alcalay RN, Gu Y, Mejia-Santana H, Cote L, Marder KS, Scarmeas N. The association between Mediterranean diet adherence and Parkinson’s disease. Mov Disord. 2012;27(6):771–4. Epub 2012/02/09. doi: 10.1002/mds.24918. PubMed PMID: 22314772; PMCID: PMC3349773.

13. Zhang X, Molsberry SA, Yeh TS, Cassidy A, Schwarzschild MA, Ascherio A, Gao X. Intake of Flavonoids and Flavonoid-Rich Foods and Mortality Risk Among Individuals With Parkinson Disease: A Prospective Cohort Study. Neurology. 2022;98(10):e1064–e76. Epub 2022/01/28. doi: 10.1212/WNL.0000000000013275. PubMed PMID: 35082171; PMCID: PMC8967390.

14. Yeh TS, Yuan C, Ascherio A, Rosner BA, Willett WC, Blacker D. Long-term Dietary Flavonoid Intake and Subjective Cognitive Decline in US Men and Women. Neurology. 2021;97(10):e1041–e56. Epub 2021/07/30. doi: 10.1212/WNL.0000000000012454. PubMed PMID: 34321362; PMCID: PMC8448553.

15. Maitra U, Conger J, Owens MMM, Ciesla L. Predicting structural features of selected flavonoids responsible for neuroprotection in a Drosophila model of Parkinson’s disease. Neurotoxicology. 2023;96:1–12. Epub 2023/02/24. doi: 10.1016/j.neuro.2023.02.008. PubMed PMID: 36822376.

16. Seigler DS, Friesen JB, Bisson J, Graham JG, Bedran-Russo A, McAlpine JB, Pauli GF. Do Certain Flavonoid IMPS Have a Vital Function? Front Nutr. 2021;8:762753. Epub 2021/12/21. doi: 10.3389/fnut.2021.762753. PubMed PMID: 34926546; PMCID: PMC8672243.

17. Bisson J, McAlpine JB, Friesen JB, Chen SN, Graham J, Pauli GF. Can Invalid Bioactives Undermine Natural Product-Based Drug Discovery? J Med Chem. 2016;59(5):1671–90. Epub 2015/10/28. doi: 10.1021/acs.jmedchem.5b01009. PubMed PMID: 26505758; PMCID: PMC4791574.

18. Maitra U, Harding T, Liang Q, Ciesla L. GardeninA confers neuroprotection against environmental toxin in a Drosophila model of Parkinson’s disease. Commun Biol. 2021;4(1):162. Epub 2021/02/07. doi: 10.1038/s42003-021-01685-2. PubMed PMID: 33547411; PMCID: PMC7864937.

19. Toppo E, Darvin SS, Esakkimuthu S, Stalin A, Balakrishna K, Sivasankaran K, Pandikumar P, Ignacimuthu S, Al-Dhabi NA. Antihyperlipidemic and hepatoprotective effects of Gardenin A in cellular and high fat diet fed rodent models. Chem Biol Interact. 2017;269:9–17. Epub 2017/03/30. doi: 10.1016/j.cbi.2017.03.013. PubMed PMID: 28351695.

20. Chiu SP, Wu MJ, Chen PY, Ho YR, Tai MH, Ho CT, Yen JH. Neurotrophic action of 5-hydroxylated polymethoxyflavones: 5-demethylnobiletin and gardenin A stimulate neuritogenesis in PC12 cells. J Agric Food Chem. 2013;61(39):9453–63. Epub 2013/09/06. doi: 10.1021/jf4024678. PubMed PMID: 24003765.

21. Alonso-Castro AJ, Gasca-Martínez D, Cortez-Mendoza LV, Alba-Betancourt C, Ruiz-Padilla AJ, Zapata-Morales JR. Evaluation of the neuropharmacological effects of Gardenin A in mice. Drug Dev Res. 2020;81(5):600–8. Epub 2020/03/18. doi: 10.1002/ddr.21659. PubMed PMID: 32181517.

22. Zhang J, Wang F, Cai W, Zhang Q, Liu Y, Li Y, Liu R, Cao G. Identification of metabolites of gardenin A in rats by combination of high-performance liquid chromatography with linear ion trap-Orbitrap mass spectrometer based on multiple data processing techniques. Biomed Chromatogr. 2015;29(3):379–87. Epub 2014/07/22. doi: 10.1002/bmc.3287. PubMed PMID: 25041995.

23. Liebisch G, Fahy E, Aoki J, Dennis EA, Durand T, Ejsing CS, Fedorova M, Feussner I, Griffiths WJ, Kofeler H, Merrill AH, Jr., Murphy RC, O’Donnell VB, Oskolkova O, Subramaniam S, Wakelam MJO, Spener F. Update on LIPID MAPS classification, nomenclature, and shorthand notation for MS-derived lipid structures. J Lipid Res. 2020;61(12):1539–55. Epub 2020/10/11. doi: 10.1194/jlr.S120001025. PubMed PMID: 33037133; PMCID: PMC7707175.

24. Vauzour D, Rendeiro C, D’Amato A, Waffo-Téguo P, Richard T, Mérillon JM, Pontifex MG, Connell E, Müller M, Butler LT, Williams CM, Spencer JPE. Anthocyanins Promote Learning through Modulation of Synaptic Plasticity Related Proteins in an Animal Model of Ageing. Antioxidants (Basel). 2021;10(8). Epub 2021/08/28. doi: 10.3390/antiox10081235. PubMed PMID: 34439483; PMCID: PMC8388918.

25. Guo Q, Kim YN, Lee BH. Protective effects of blueberry drink on cognitive impairment induced by chronic mild stress in adult rats. Nutr Res Pract. 2017;11(1):25–32. Epub 2017/02/15. doi: 10.4162/nrp.2017.11.1.25. PubMed PMID: 28194262; PMCID: PMC5300943.

26. Papandreou MA, Dimakopoulou A, Linardaki ZI, Cordopatis P, Klimis-Zacas D, Margarity M, Lamari FN. Effect of a polyphenol-rich wild blueberry extract on cognitive performance of mice, brain antioxidant markers and acetylcholinesterase activity. Behav Brain Res. 2009;198(2):352–8. Epub 2008/12/06. doi: 10.1016/j.bbr.2008.11.013. PubMed PMID: 19056430.

27. Dodd GF, Williams CM, Butler LT, Spencer JPE. Acute effects of flavonoid-rich blueberry on cognitive and vascular function in healthy older adults. Nutrition and Healthy Aging. 2019;5(2):119–32.

28. Krikorian R, Shidler MD, Nash TA, Kalt W, Vinqvist-Tymchuk MR, Shukitt-Hale B, Joseph JA. Blueberry supplementation improves memory in older adults. J Agric Food Chem. 2010;58(7):3996–4000. Epub 2010/01/06. doi: 10.1021/jf9029332. PubMed PMID: 20047325; PMCID: PMC2850944.

29. Nan S, Wang P, Zhang Y, Fan J. Epigallocatechin-3-Gallate Provides Protection Against Alzheimer’s Disease-Induced Learning and Memory Impairments in Rats. Drug Des Devel Ther. 2021;15:2013–24. Epub 2021/05/21. doi: 10.2147/DDDT.S289473. PubMed PMID: 34012254; PMCID: PMC8128347.

30. Wei BB, Liu MY, Zhong X, Yao WF, Wei MJ. Increased BBB permeability contributes to EGCG-caused cognitive function improvement in natural aging rats: pharmacokinetic and distribution analyses. Acta Pharmacol Sin. 2019;40(11):1490–500. Epub 2019/05/17. doi: 10.1038/s41401-019-0243-7. PubMed PMID: 31092885; PMCID: PMC6888860.

31. Ahn JW, Kim S, Ko S, Kim YH, Jeong JH, Chung S. Modified (-)-gallocatechin gallate-enriched green tea extract rescues age-related cognitive deficits by restoring hippocampal synaptic plasticity. Biochem Biophys Rep. 2022;29:101201. Epub 2022/02/25. doi: 10.1016/j.bbrep.2022.101201. PubMed PMID: 35198737; PMCID: PMC8841891.

32. Scholey A, Downey LA, Ciorciari J, Pipingas A, Nolidin K, Finn M, Wines M, Catchlove S, Terrens A, Barlow E, Gordon L, Stough C. Acute neurocognitive effects of epigallocatechin gallate (EGCG). Appetite. 2012;58(2):767–70. Epub 2011/12/01. doi: 10.1016/j.appet.2011.11.016. PubMed PMID: 22127270.

33. Mastroiacovo D, Kwik-Uribe C, Grassi D, Necozione S, Raffaele A, Pistacchio L, Righetti R, Bocale R, Lechiara MC, Marini C, Ferri C, Desideri G. Cocoa flavanol consumption improves cognitive function, blood pressure control, and metabolic profile in elderly subjects: the Cocoa, Cognition, and Aging (CoCoA) Study--a randomized controlled trial. Am J Clin Nutr. 2015;101(3):538–48. Epub 2015/03/04. doi: 10.3945/ajcn.114.092189. PubMed PMID: 25733639; PMCID: PMC4340060.

34. Desideri G, Kwik-Uribe C, Grassi D, Necozione S, Ghiadoni L, Mastroiacovo D, Raffaele A, Ferri L, Bocale R, Lechiara MC, Marini C, Ferri C. Benefits in cognitive function, blood pressure, and insulin resistance through cocoa flavanol consumption in elderly subjects with mild cognitive impairment: the Cocoa, Cognition, and Aging (CoCoA) study. Hypertension. 2012;60(3):794–801. Epub 2012/08/16. doi: 10.1161/HYPERTENSIONAHA.112.193060. PubMed PMID: 22892813.

35. Bisson JF, Nejdi A, Rozan P, Hidalgo S, Lalonde R, Messaoudi M. Effects of long-term administration of a cocoa polyphenolic extract (Acticoa powder) on cognitive performances in aged rats. Br J Nutr. 2008;100(1):94–101. Epub 2008/01/09. doi: 10.1017/S0007114507886375. PubMed PMID: 18179729.

36. Rozan P, Hidalgo S, Nejdi A, Bisson JF, Lalonde R, Messaoudi M. Preventive antioxidant effects of cocoa polyphenolic extract on free radical production and cognitive performances after heat exposure in Wistar rats. J Food Sci. 2007;72(3):S203–6. Epub 2007/11/13. doi: 10.1111/j.1750-3841.2007.00297.x. PubMed PMID: 17995815.

37. Sashourpour M, Zahri S, Radjabian T, Ruf V, Pan-Montojo F, Morshedi D. A study on the modulation of alpha-synuclein fibrillation by Scutellaria pinnatifida extracts and its neuroprotective properties. PLoS One. 2017;12(9):e0184483. Epub 2017/09/29. doi: 10.1371/journal.pone.0184483. PubMed PMID: 28957336; PMCID: PMC5619708.

38. Chen M, Wang T, Yue F, Li X, Wang P, Li Y, Chan P, Yu S. Tea polyphenols alleviate motor impairments, dopaminergic neuronal injury, and cerebral α-synuclein aggregation in MPTP-intoxicated parkinsonian monkeys. Neuroscience. 2015;286:383–92. Epub 2014/12/17. doi: 10.1016/j.neuroscience.2014.12.003. PubMed PMID: 25498223.

39. Tikhonova MA, Tikhonova NG, Tenditnik MV, Ovsyukova MV, Akopyan AA, Dubrovina NI, Amstislavskaya TG, Khlestkina EK. Effects of Grape Polyphenols on the Life Span and Neuroinflammatory Alterations Related to Neurodegenerative Parkinson Disease-Like Disturbances in Mice. Molecules. 2020;25(22). Epub 2020/11/20. doi: 10.3390/molecules25225339. PubMed PMID: 33207644; PMCID: PMC7696792.

40. Zhang Y, Wu Q, Zhang L, Wang Q, Yang Z, Liu J, Feng L. Caffeic acid reduces A53T alpha-synuclein by activating JNK/Bcl-2-mediated autophagy in vitro and improves behaviour and protects dopaminergic neurons in a mouse model of Parkinson’s disease. Pharmacol Res. 2019;150:104538. Epub 2019/11/11. doi: 10.1016/j.phrs.2019.104538. PubMed PMID: 31707034.

41. Kuang S, Yang L, Rao Z, Zhong Z, Li J, Zhong H, Dai L, Tang X. Effects of Ginkgo Biloba Extract on A53T alpha-Synuclein Transgenic Mouse Models of Parkinson’s Disease. Can J Neurol Sci. 2018;45(2):182–7. Epub 2018/03/07. doi: 10.1017/cjn.2017.268. PubMed PMID: 29506601.

42. Blesa J, Trigo-Damas I, Quiroga-Varela A, Jackson-Lewis VR. Oxidative stress and Parkinson’s disease. Front Neuroanat. 2015;9:91. Epub 2015/07/29. doi: 10.3389/fnana.2015.00091. PubMed PMID: 26217195; PMCID: PMC4495335.

43. Zhang LF, Yu XL, Ji M, Liu SY, Wu XL, Wang YJ, Liu RT. Resveratrol alleviates motor and cognitive deficits and neuropathology in the A53T alpha-synuclein mouse model of Parkinson’s disease. Food Funct. 2018;9(12):6414–26. Epub 2018/11/22. doi: 10.1039/c8fo00964c. PubMed PMID: 30462117.

44. He XJ, Uchida K, Megumi C, Tsuge N, Nakayama H. Dietary curcumin supplementation attenuates 1-methyl-4-phenyl-1,2,3,6-tetrahydropyridine (MPTP) neurotoxicity in C57BL mice. J Toxicol Pathol. 2015;28(4):197-206. Epub 2015/11/06. doi: 10.1293/tox.2015-0020. PubMed PMID: 26538809; PMCID: PMC4604129.

45. Zhu H, Zhang H, Hou B, Xu B, Ji L, Wu Y. Curcumin Regulates Gut Microbiota and Exerts a Neuroprotective Effect in the MPTP Model of Parkinson’s Disease. Evid Based Complement Alternat Med. 2022;2022:9110560. Epub 2022/12/06. doi: 10.1155/2022/9110560. PubMed PMID: 36467550; PMCID: PMC9715342 publication of this paper.

46. Jain SK, Rains J, Croad J, Larson B, Jones K. Curcumin supplementation lowers TNF-alpha, IL-6, IL-8, and MCP-1 secretion in high glucose-treated cultured monocytes and blood levels of TNF-alpha, IL-6, MCP-1, glucose, and glycosylated hemoglobin in diabetic rats. Antioxid Redox Signal. 2009;11(2):241-9. Epub 2008/11/04. doi: 10.1089/ars.2008.2140. PubMed PMID: 18976114; PMCID: PMC2933148.

47. Zhou T, Zhu M, Liang Z. (-)-Epigallocatechin-3-gallate modulates peripheral immunity in the MPTP-induced mouse model of Parkinson’s disease. Mol Med Rep. 2018;17(4):4883–8. Epub 2018/01/25. doi: 10.3892/mmr.2018.8470. PubMed PMID: 29363729; PMCID: PMC5865947.

48. Kim SR, Seong KJ, Kim WJ, Jung JY. Epigallocatechin Gallate Protects against Hypoxia-Induced Inflammation in Microglia via NF-κB Suppression and Nrf-2/HO-1 Activation. Int J Mol Sci. 2022;23(7). Epub 2022/04/13. doi: 10.3390/ijms23074004. PubMed PMID: 35409364; PMCID: PMC8999549.

49. Nagayoshi H, Murayama N, Takenaka S, Kim V, Kim D, Komori M, Yamazaki H, Guengerich FP, Shimada T. Roles of cytochrome P450 2A6 in the oxidation of flavone, 4’-hydroxyflavone, and 4’-, 3’-, and 2’-methoxyflavones by human liver microsomes. Xenobiotica. 2021;51(9):995-1009. Epub 2021/07/06. doi: 10.1080/00498254.2021.1950866. PubMed PMID: 34224301; PMCID: PMC8719450.

50. Abdelazeem KNM, Kalo MZ, Beer-Hammer S, Lang F. The gut microbiota metabolite urolithin A inhibits NF-kappaB activation in LPS stimulated BMDMs. Sci Rep. 2021;11(1):7117. Epub 2021/03/31. doi: 10.1038/s41598-021-86514-6. PubMed PMID: 33782464; PMCID: PMC8007722.

51. Melrose J. The Potential of Flavonoids and Flavonoid Metabolites in the Treatment of Neurodegenerative Pathology in Disorders of Cognitive Decline. Antioxidants (Basel). 2023;12(3). Epub 2023/03/30. doi: 10.3390/antiox12030663. PubMed PMID: 36978911; PMCID: PMC10045397.

52. Paraiso IL, Revel JS, Choi J, Miranda CL, Lak P, Kioussi C, Bobe G, Gombart AF, Raber J, Maier CS, Stevens JF. Targeting the Liver-Brain Axis with Hop-Derived Flavonoids Improves Lipid Metabolism and Cognitive Performance in Mice. Mol Nutr Food Res. 2020;64(15):e2000341. Epub 2020/07/07. doi: 10.1002/mnfr.202000341. PubMed PMID: 32627931; PMCID: PMC8693899.

53. Cataldi S, Arcuri C, Hunot S, Legeron FP, Mecca C, Garcia-Gil M, Lazzarini A, Codini M, Beccari T, Tasegian A, Fioretti B, Traina G, Ambesi-Impiombato FS, Curcio F, Albi E. Neutral Sphingomyelinase Behaviour in Hippocampus Neuroinflammation of MPTP-Induced Mouse Model of Parkinson’s Disease and in Embryonic Hippocampal Cells. Mediators Inflamm. 2017;2017:2470950. Epub 2018/01/19. doi: 10.1155/2017/2470950. PubMed PMID: 29343884; PMCID: PMC5733979.

54. Kettwig M, Klemp H, Nessler S, Streit F, Kratzner R, Rosewich H, Gartner J. Targeted metabolomics revealed changes in phospholipids during the development of neuroinflammation in Abcd1(tm1Kds) mice and X-linked adrenoleukodystrophy patients. J Inherit Metab Dis. 2021;44(5):1174–85. Epub 2021/04/16. doi: 10.1002/jimd.12389. PubMed PMID: 33855724.

55. Karonen M. Insights into Polyphenol-Lipid Interactions: Chemical Methods, Molecular Aspects and Their Effects on Membrane Structures. Plants (Basel). 2022;11(14). Epub 2022/07/28. doi: 10.3390/plants11141809. PubMed PMID: 35890443; PMCID: PMC9317924.

56. Goldberg NR, Hampton T, McCue S, Kale A, Meshul CK. Profiling changes in gait dynamics resulting from progressive 1-methyl-4-phenyl-1,2,3,6-tetrahydropyridine-induced nigrostriatal lesioning. J Neurosci Res. 2011;89(10):1698-706. Epub 2011/07/13. doi: 10.1002/jnr.22699. PubMed PMID: 21748776.

57. Quinn JF, Kelly MJ, Harris CJ, Hack W, Gray NE, Kulik V, Bostick Z, Brumbach BH, Copenhaver PF. The novel estrogen receptor modulator STX attenuates Amyloid-beta neurotoxicity in the 5XFAD mouse model of Alzheimer’s disease. Neurobiol Dis. 2022;174:105888. Epub 2022/10/10. doi: 10.1016/j.nbd.2022.105888. PubMed PMID: 36209948.

