## Supplementary materials for "Gardenin A improves cognitive and motor function in A53T-α-syn mice"

SUPPLEMENTARY FIGURES

**GA bioavailability in A53TSyn mouse brain tissues.**

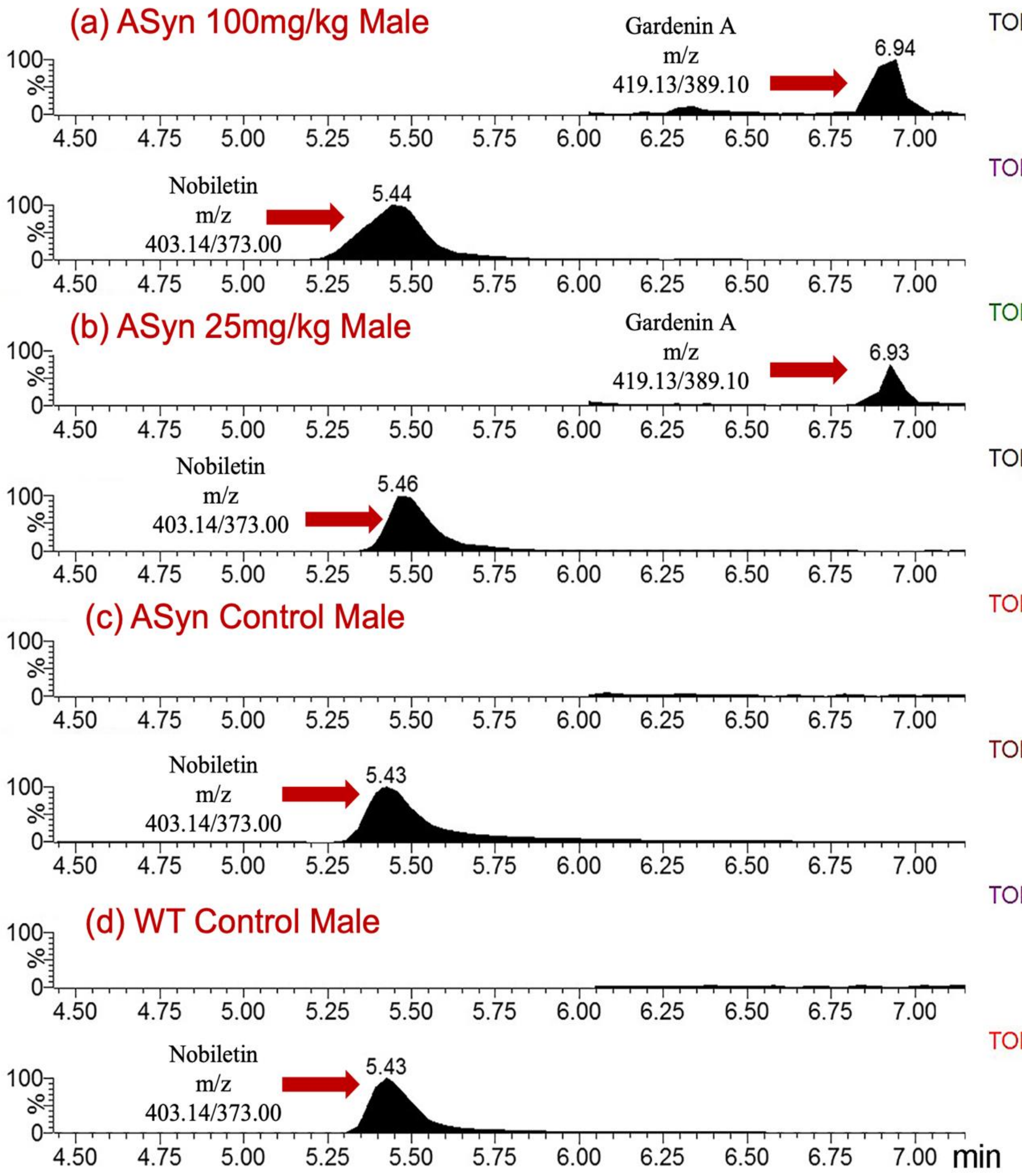


**Figure S1:** Example chromatograms presenting nobiletin and GA peaks in the brain sample obtained from a male mouse treated with (a) 100 mg/kg GA and (b) 25 mg/kg GA. Example chromatograms presenting the peak of nobiletin and the absence on GA peak in the brain sample obtained from a (c) A53TSyn male mouse and (d) WT male mouse treated with a vehicle.

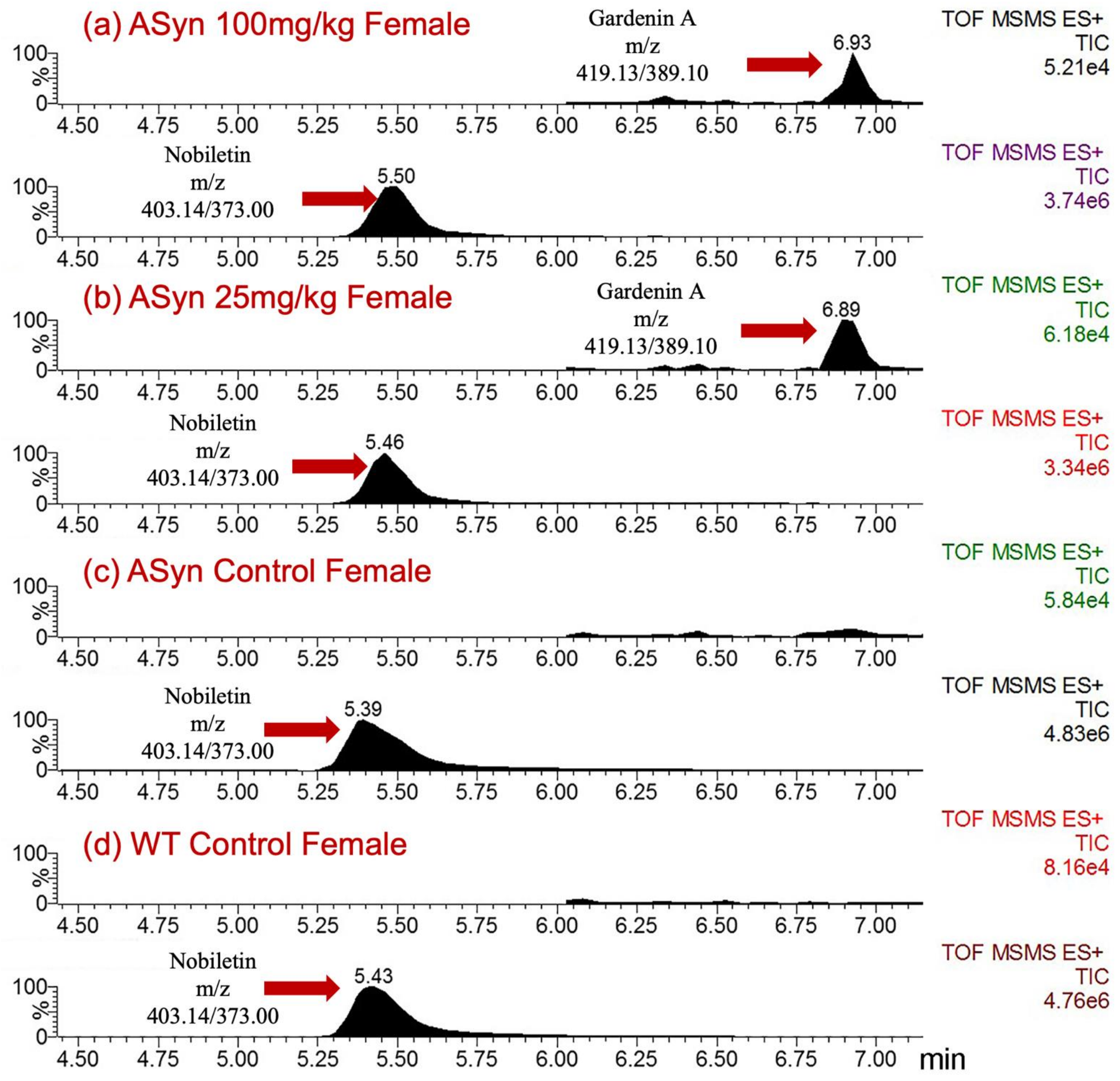


**Figure S2:** Example chromatograms presenting nobiletin and GA peaks in the brain sample obtained from a female mouse treated with (a) 100 mg/kg GA and (b) 25 mg/kg GA. Example chromatograms presenting the peak of nobiletin and the absence on GA peak in the brain sample obtained from a (c) A53TSyn female mouse and (d) WT female mouse treated with a vehicle.

Represented below are the Total ion chromatograms for each individual sample (male/female) for both GA treated (100/25 mg/kg) and vehicle treated (A53TSyn/WT control) mice. GA peaks were consistently across all the individual mice samples. Nobiletin (internal standard) could also be recovered from all the individual mice brain samples.

**GarA_Sample1_Vehicle WT Control male**

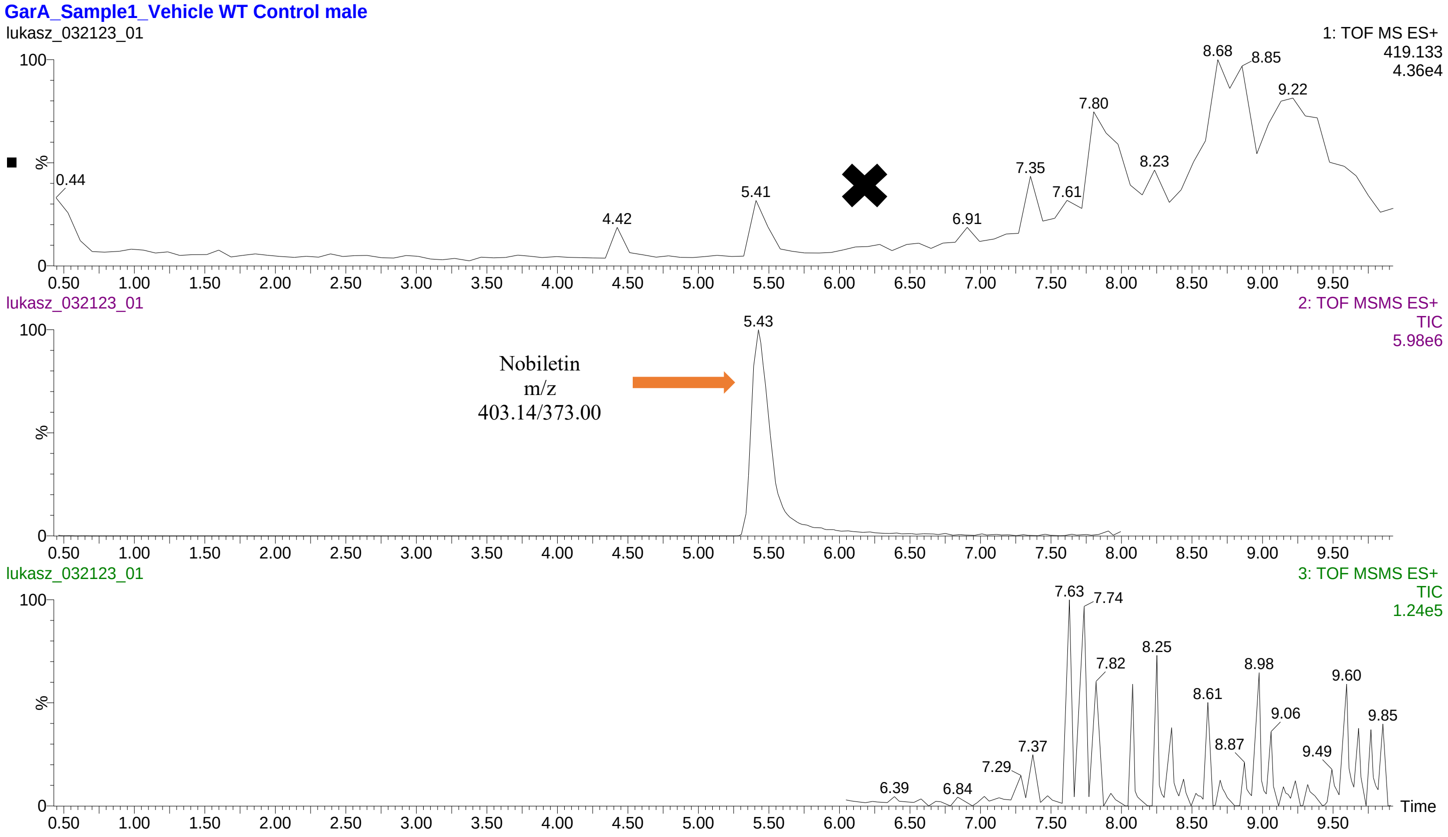


**GarA_Sample1_Vehicle aSyn Control male**

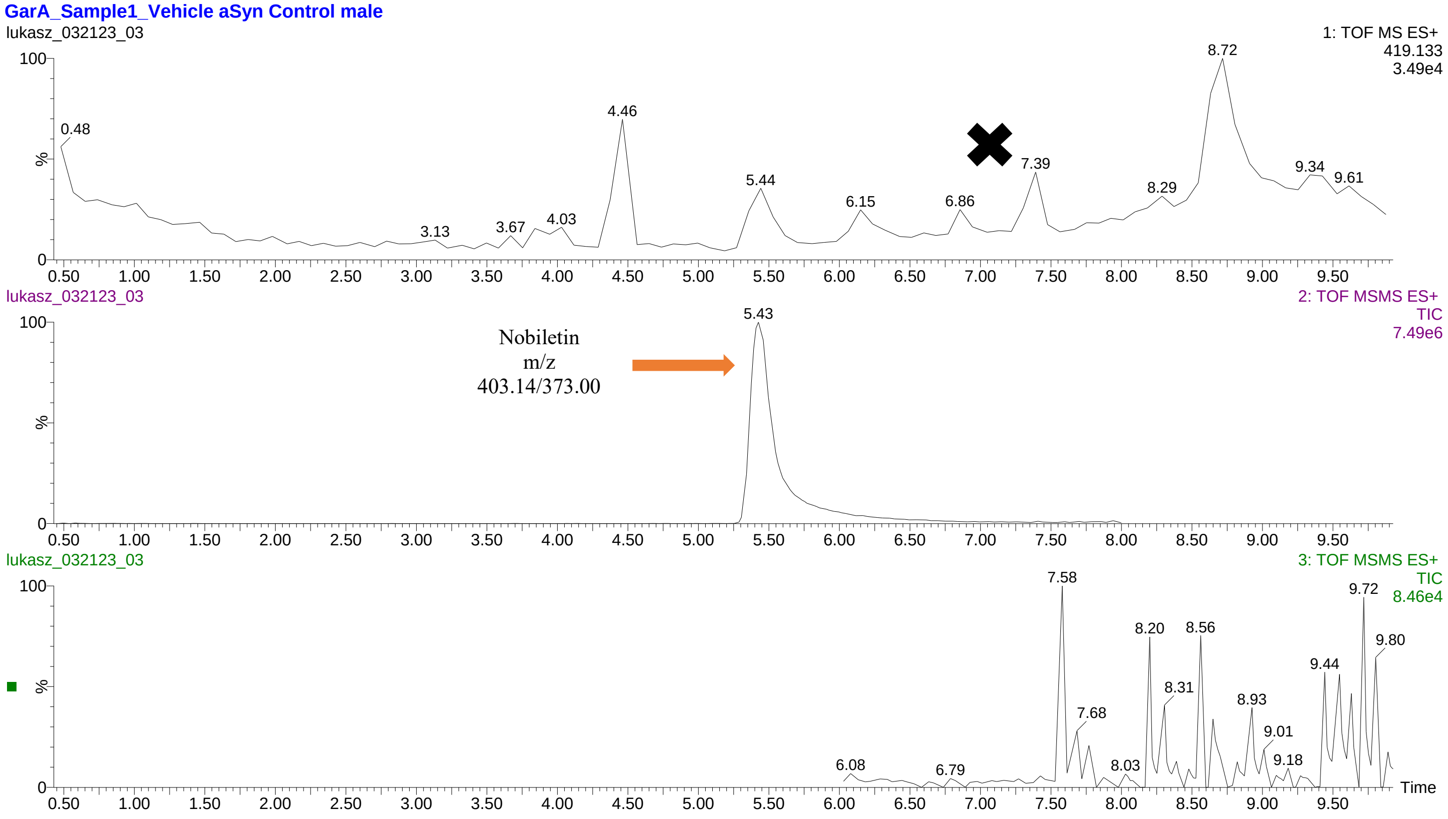

**GarA_Sample1_Vehicle WT Control female**

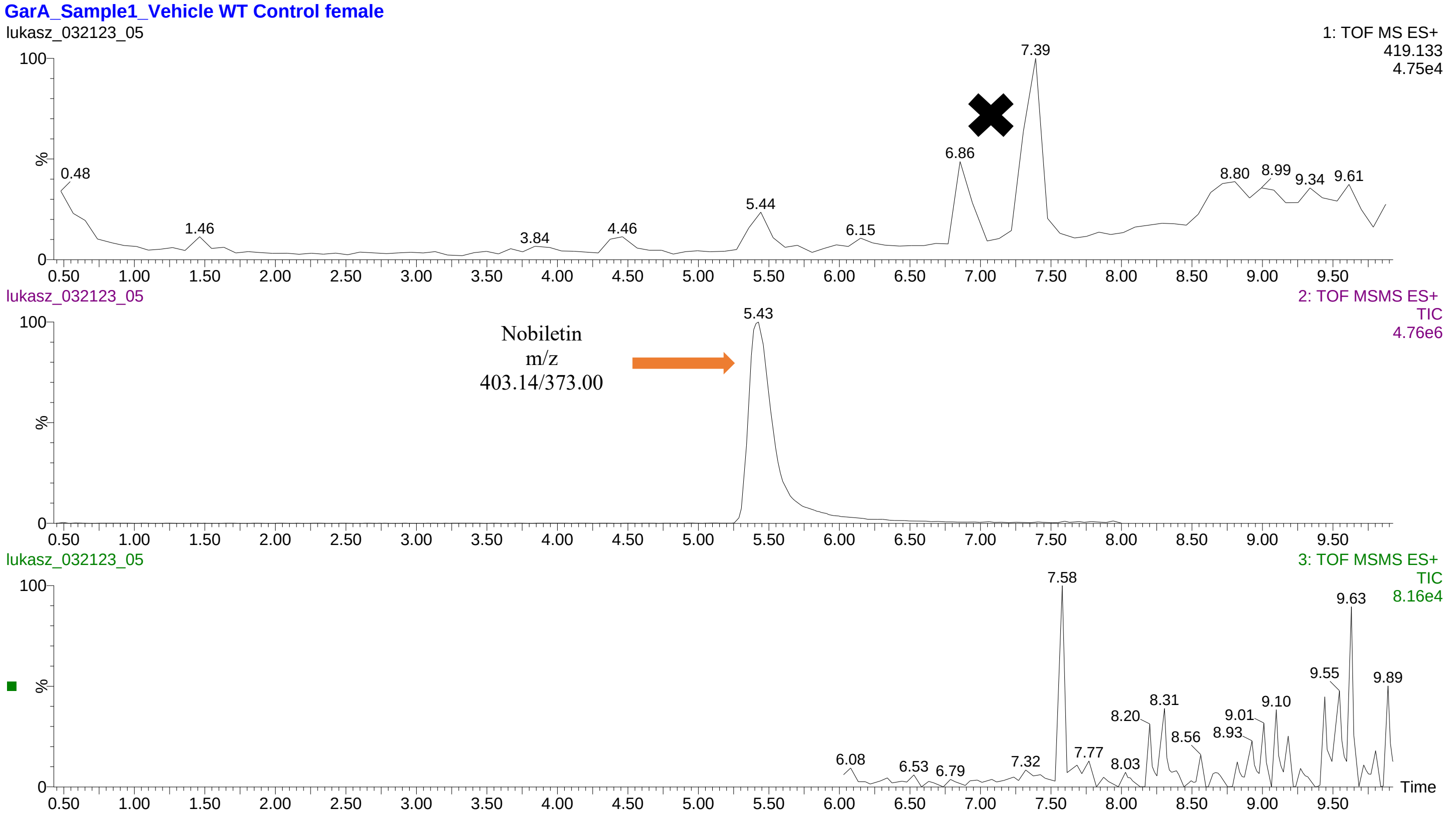


**GarA_Sample1_Vehicle aSyn Control female**

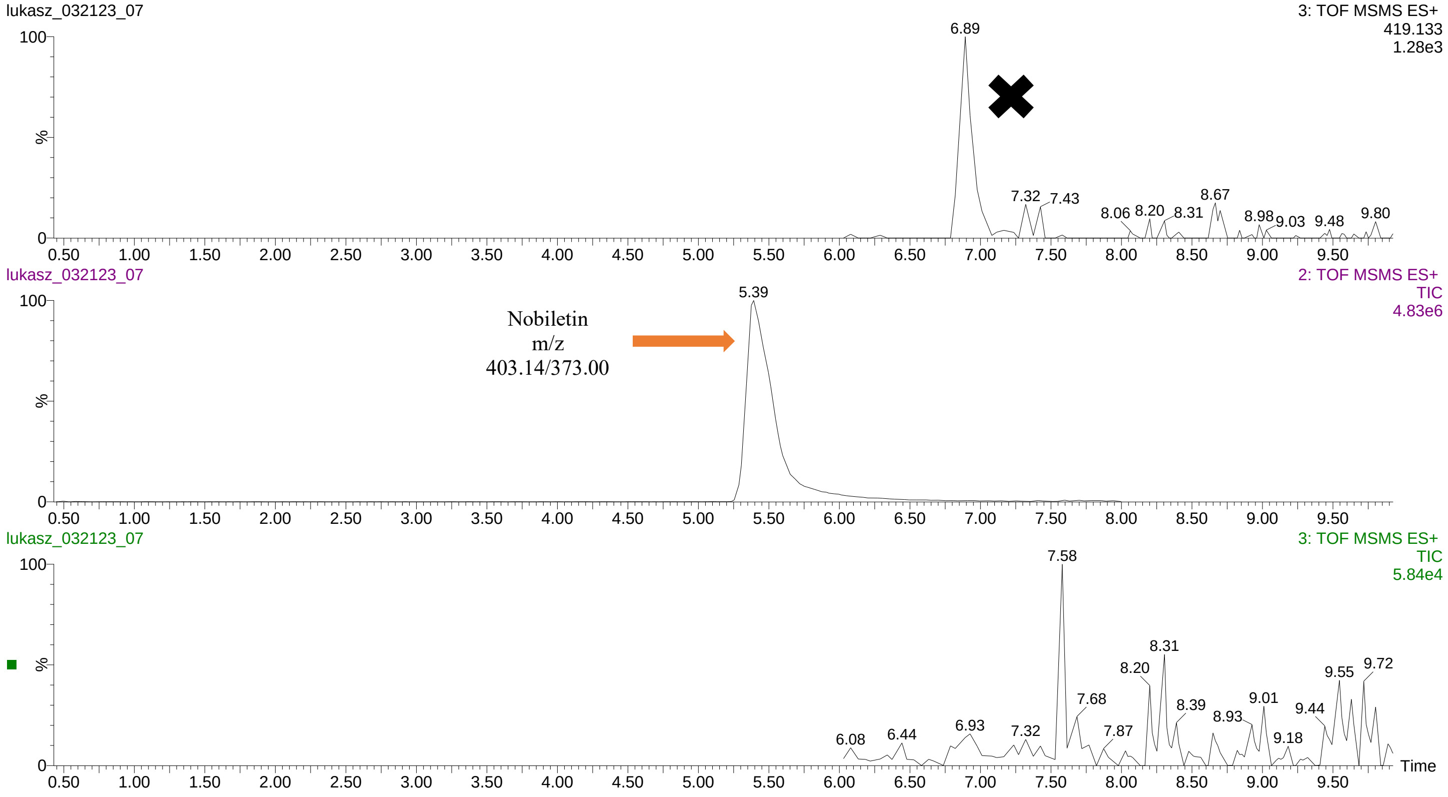

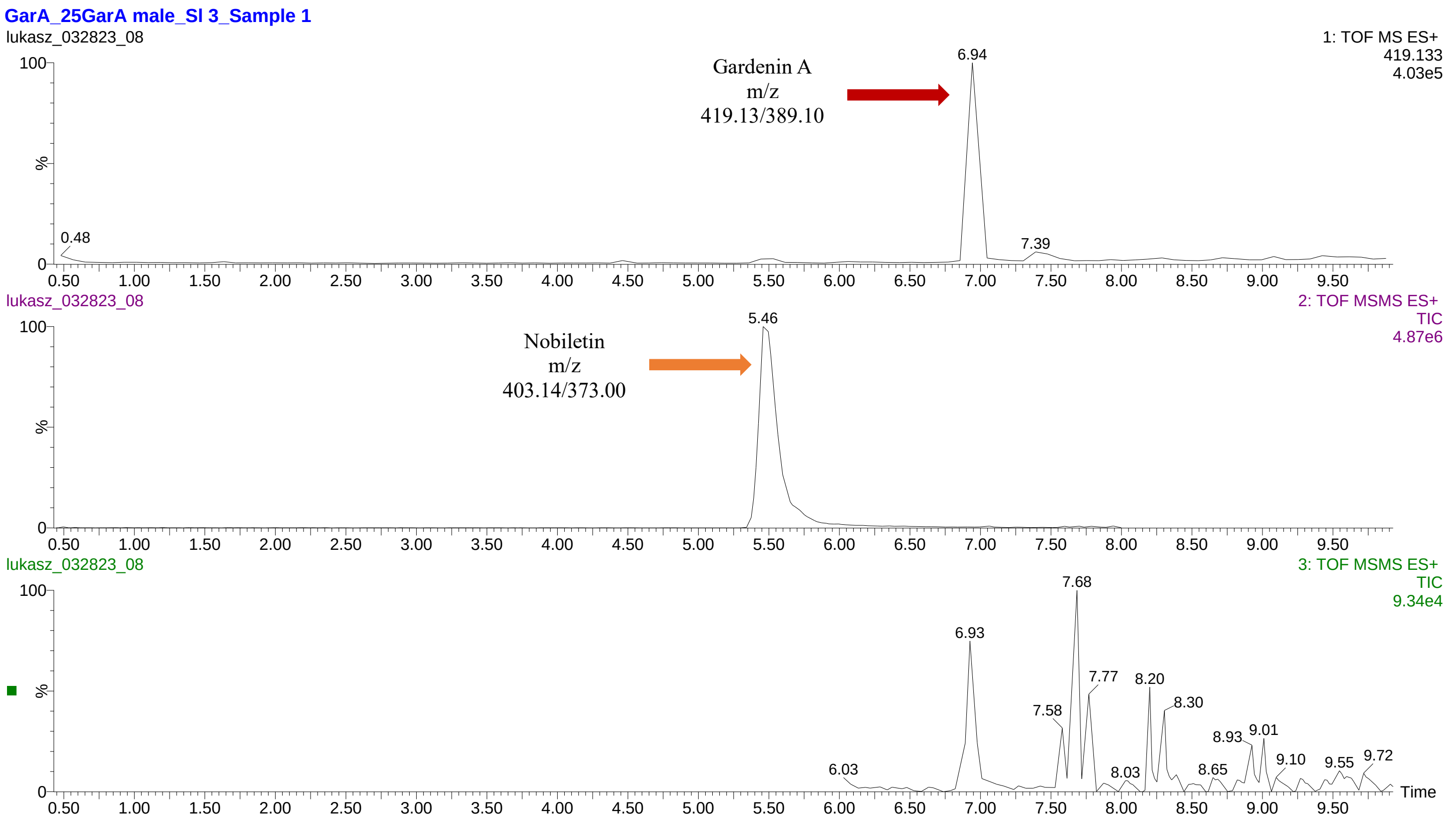
GarA_25GarA male_SI 3_Sample 1
lukasz_032823_08
1: TOF MS ES+
419.133
4.03e5
Gardenin A
m/z
419.13/389.10
6.94
0.48
7.39
2: TOF MSMS ES+
TIC
4.87e6
Nobiletin
m/z
403.14/373.00
5.46
3: TOF MSMS ES+
TIC
9.34e4
7.68
6.93
7.77
8.20
7.58
8.30
8.93
9.01
6.03
8.03
8.65
9.10
9.55
9.72
Time

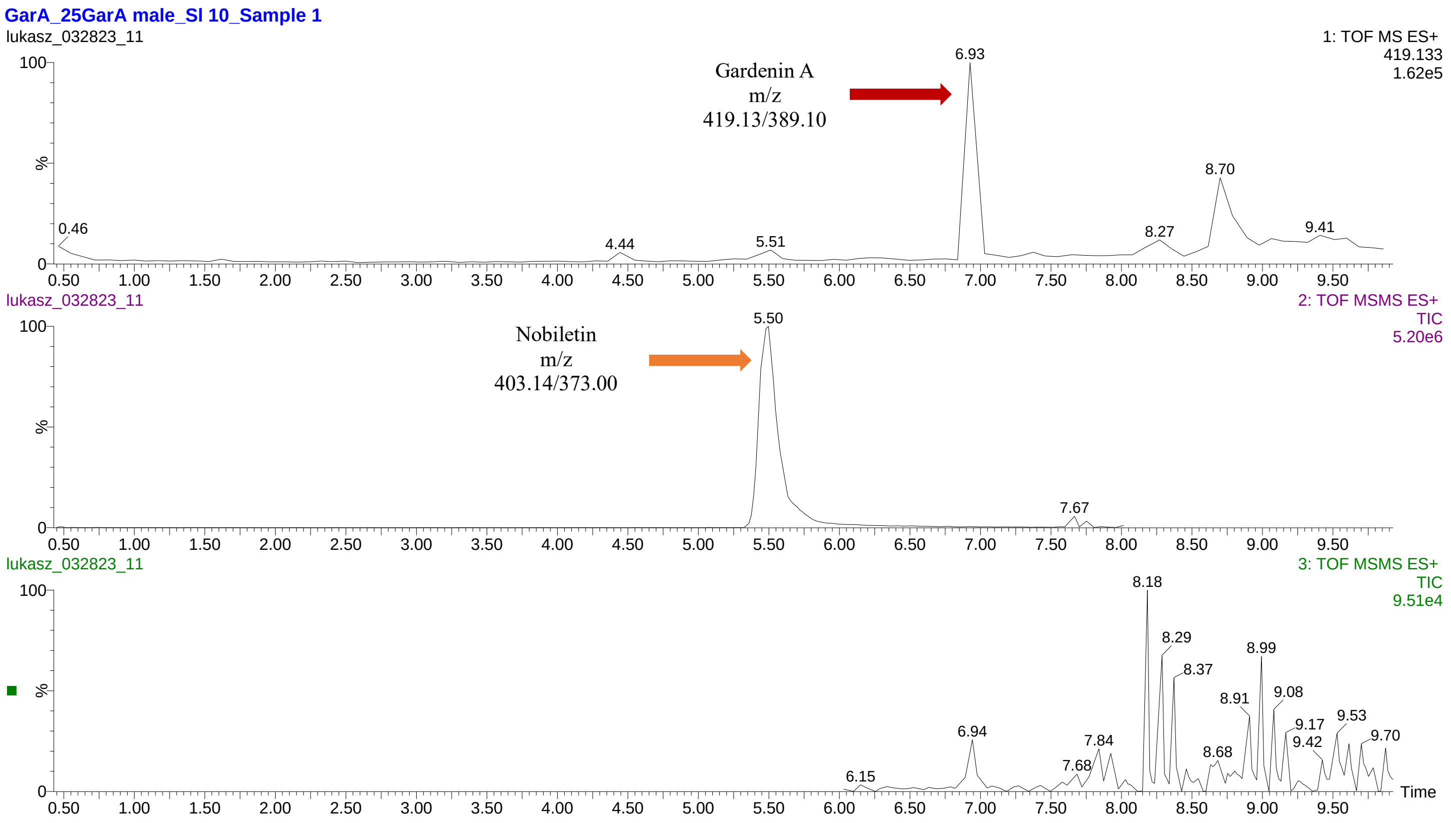
GarA_25GarA male_SI 10_Sample 1
lukasz_032823_11
1: TOF MS ES+
419.133
1.62e5
Gardenin A
m/z
419.13/389.10
6.93
8.70
0.46
4.44
5.51
8.27
9.41
2: TOF MSMS ES+
TIC
5.20e6
Nobiletin
m/z
403.14/373.00
5.50
7.67
3: TOF MSMS ES+
TIC
9.51e4
8.18
8.29
8.99
8.37
8.91
9.08
9.17
9.42
9.53
9.70
6.94
7.84
7.68
8.68
6.15
Time

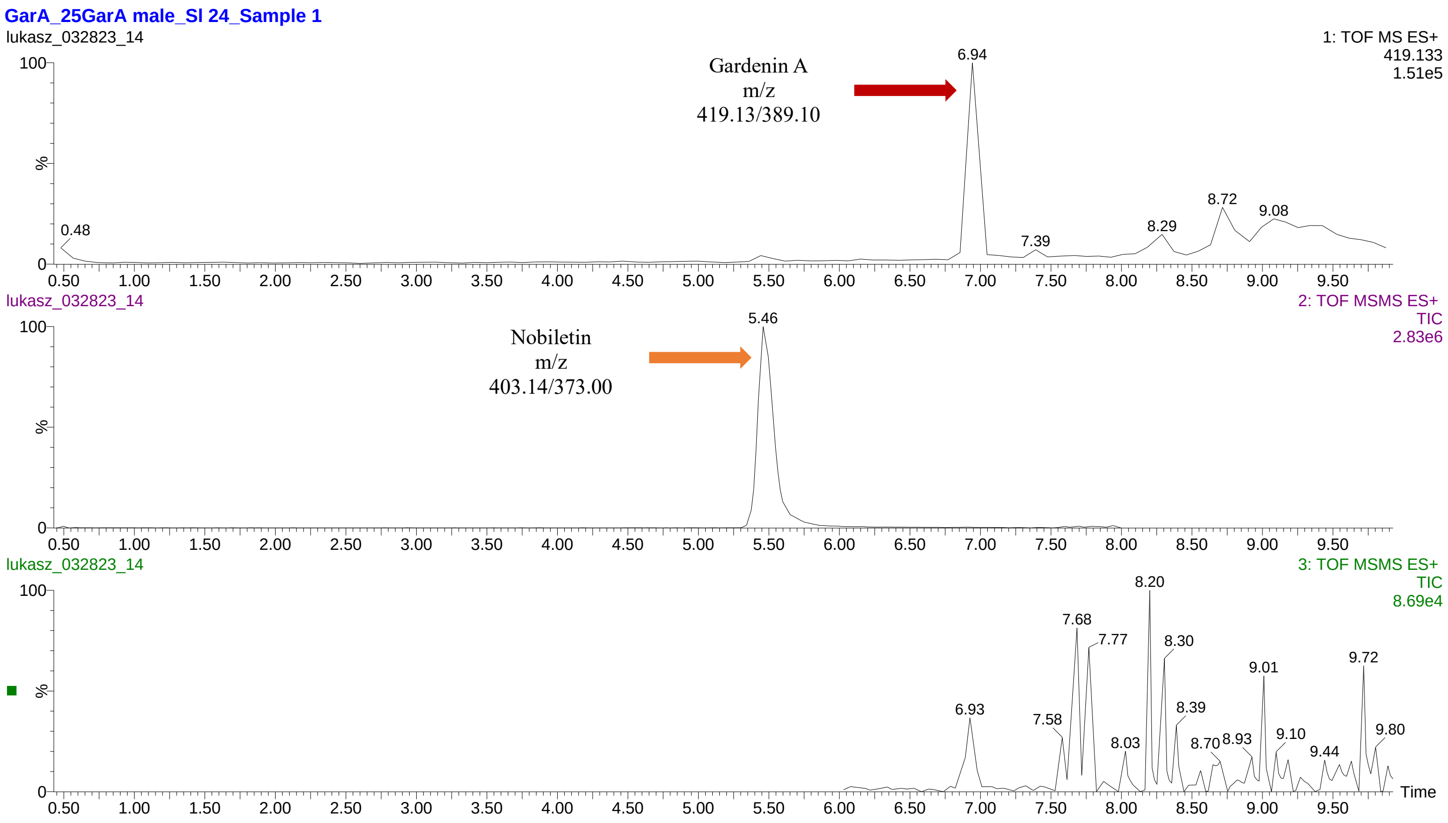

GarA_25GarA male_SI 24_Sample 1
lukasz_032823_14
1: TOF MS ES+
419.133
1.51e5
Gardenin A
m/z
419.13/389.10
6.94
0.48
7.39
8.29
8.72
9.08
2: TOF MSMS ES+
TIC
2.83e6
Nobiletin
m/z
403.14/373.00
5.46
3: TOF MSMS ES+
TIC
8.69e4
8.20
7.68
7.77
8.30
9.01
9.72
6.93
7.58
8.39
8.03
8.70
8.93
9.10
9.44
9.80
%
Time


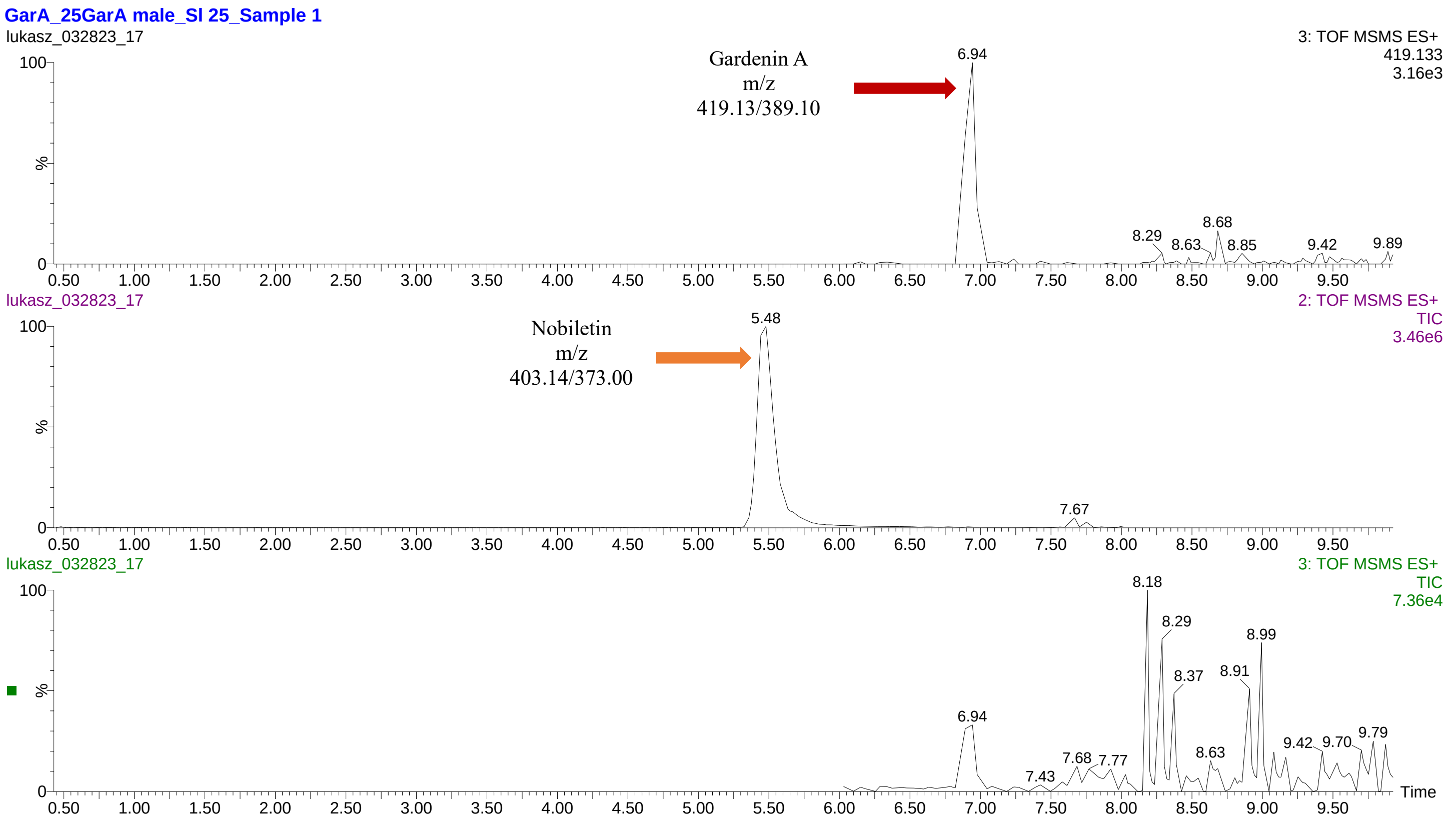

GarA_25GarA male_SI 25_Sample 1
lukasz_032823_17
3: TOF MSMS ES+
419.133
3.16e3
Gardenin A
m/z
419.13/389.10
6.94
8.29
8.63
8.68
8.85
9.42
9.89
2: TOF MSMS ES+
TIC
3.46e6
Nobiletin
m/z
403.14/373.00
5.48
7.67
3: TOF MSMS ES+
TIC
7.36e4
8.18
8.29
8.99
8.37
8.91
6.94
7.43
7.68
7.77
8.63
9.42
9.70
9.79
%
Time

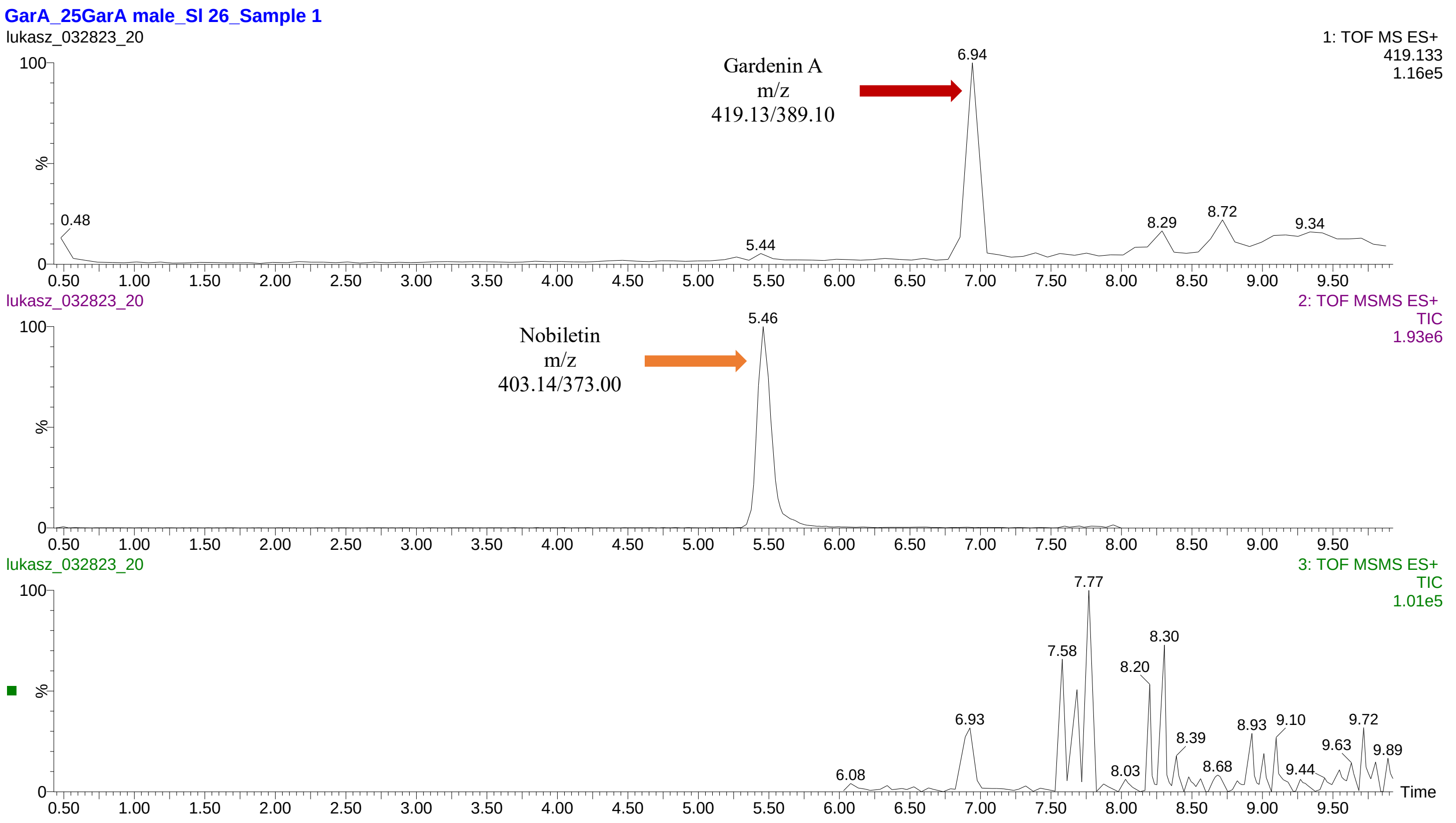
GarA_25GarA male_SI 26_Sample 1
lukasz_032823_20
1: TOF MS ES+
419.133
1.16e5
Gardenin A
m/z
419.13/389.10
6.94
0.48
5.44
8.29
8.72
9.34
lukasz_032823_20
2: TOF MSMS ES+
TIC
1.93e6
Nobiletin
m/z
403.14/373.00
5.46
lukasz_032823_20
3: TOF MSMS ES+
TIC
1.01e5
7.77
8.30
7.58
8.20
6.93
8.39
8.93
9.10
9.72
9.63
9.89
6.08
8.03
8.68
9.44
%
Time

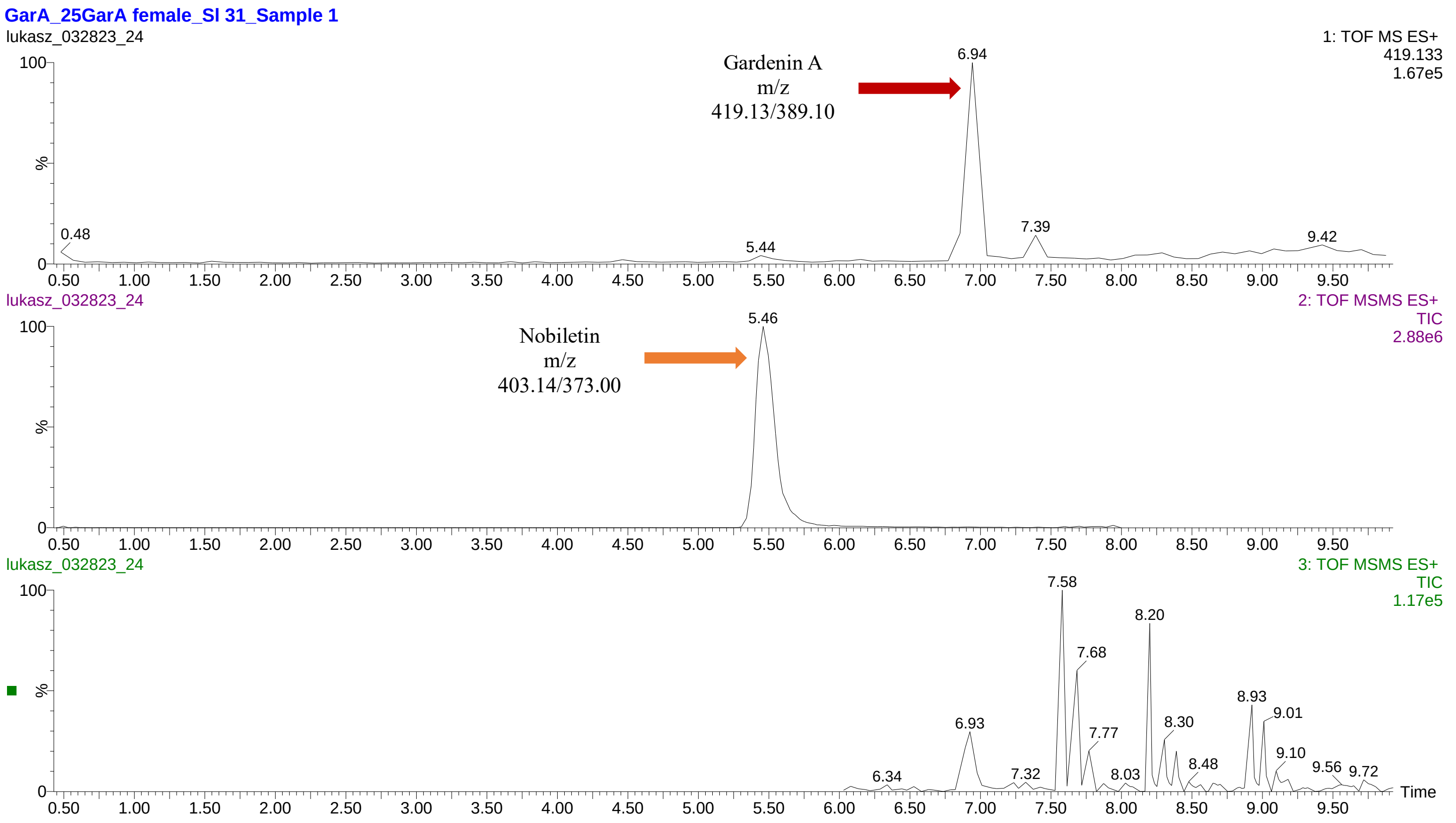
GarA_25GarA female_SI 31_Sample 1
lukasz_032823_24
1: TOF MS ES+
419.133
1.67e5
Gardenin A
m/z
419.13/389.10
6.94
0.48
5.44
7.39
9.42
lukasz_032823_24
2: TOF MSMS ES+
TIC
2.88e6
Nobiletin
m/z
403.14/373.00
5.46
lukasz_032823_24
3: TOF MSMS ES+
TIC
1.17e5
7.58
8.20
7.68
8.93
9.01
6.93
8.30
7.77
9.10
8.48
9.56
9.72
6.34
7.32
8.03
%
Time

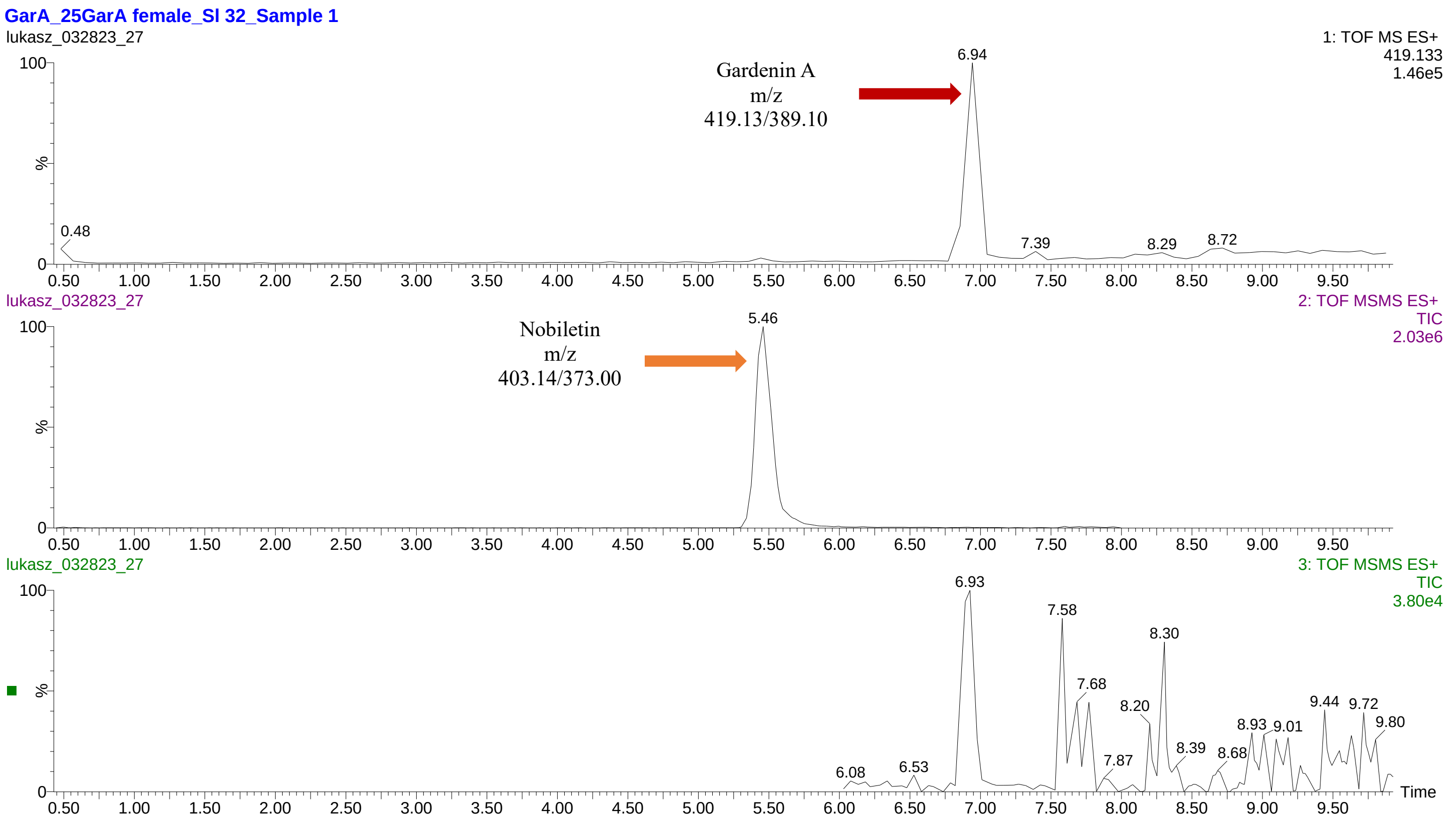
GarA_25GarA female_SI 32_Sample 1
lukasz_032823_27
1: TOF MS ES+
419.133
1.46e5
Gardenin A
m/z
419.13/389.10
6.94
0.48
7.39
8.29
8.72
%
lukasz_032823_27
2: TOF MSMS ES+
TIC
2.03e6
Nobiletin
m/z
403.14/373.00
5.46
lukasz_032823_27
3: TOF MSMS ES+
TIC
3.80e4
6.93
7.58
8.30
7.68
8.20
9.44
9.72
9.80
8.93
9.01
8.39
8.68
7.87
6.08
6.53
Time

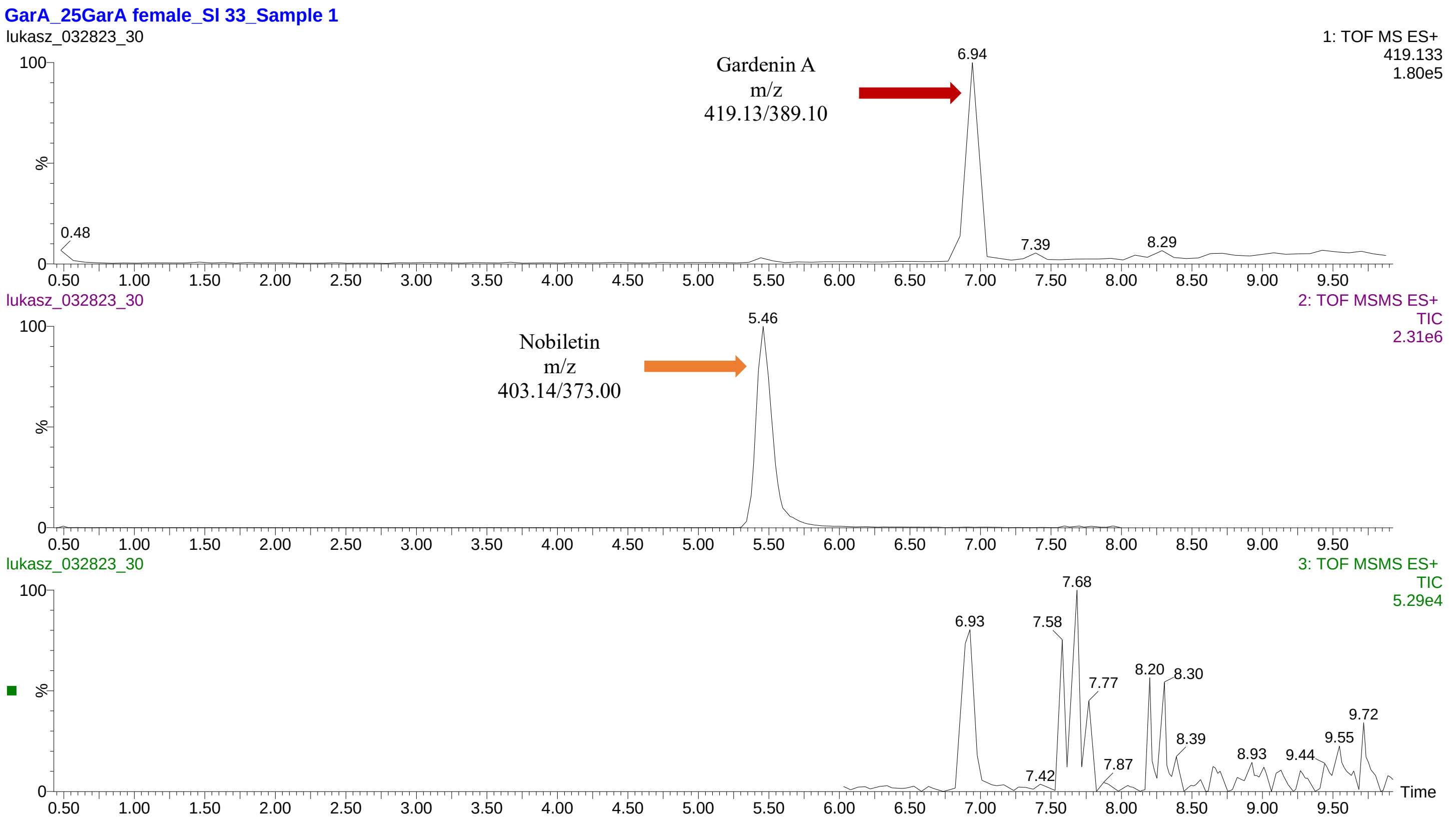
GarA_25GarA female_SI 33_Sample 1
lukasz_032823_30
1: TOF MS ES+
419.133
1.80e5
Gardenin A
m/z
419.13/389.10
6.94
0.48
7.39
8.29
%
lukasz_032823_30
2: TOF MSMS ES+
TIC
2.31e6
Nobiletin
m/z
403.14/373.00
5.46
lukasz_032823_30
3: TOF MSMS ES+
TIC
5.29e4
7.68
6.93
7.58
7.77
8.20
8.30
9.72
8.39
9.55
8.93
9.44
7.87
7.42
Time

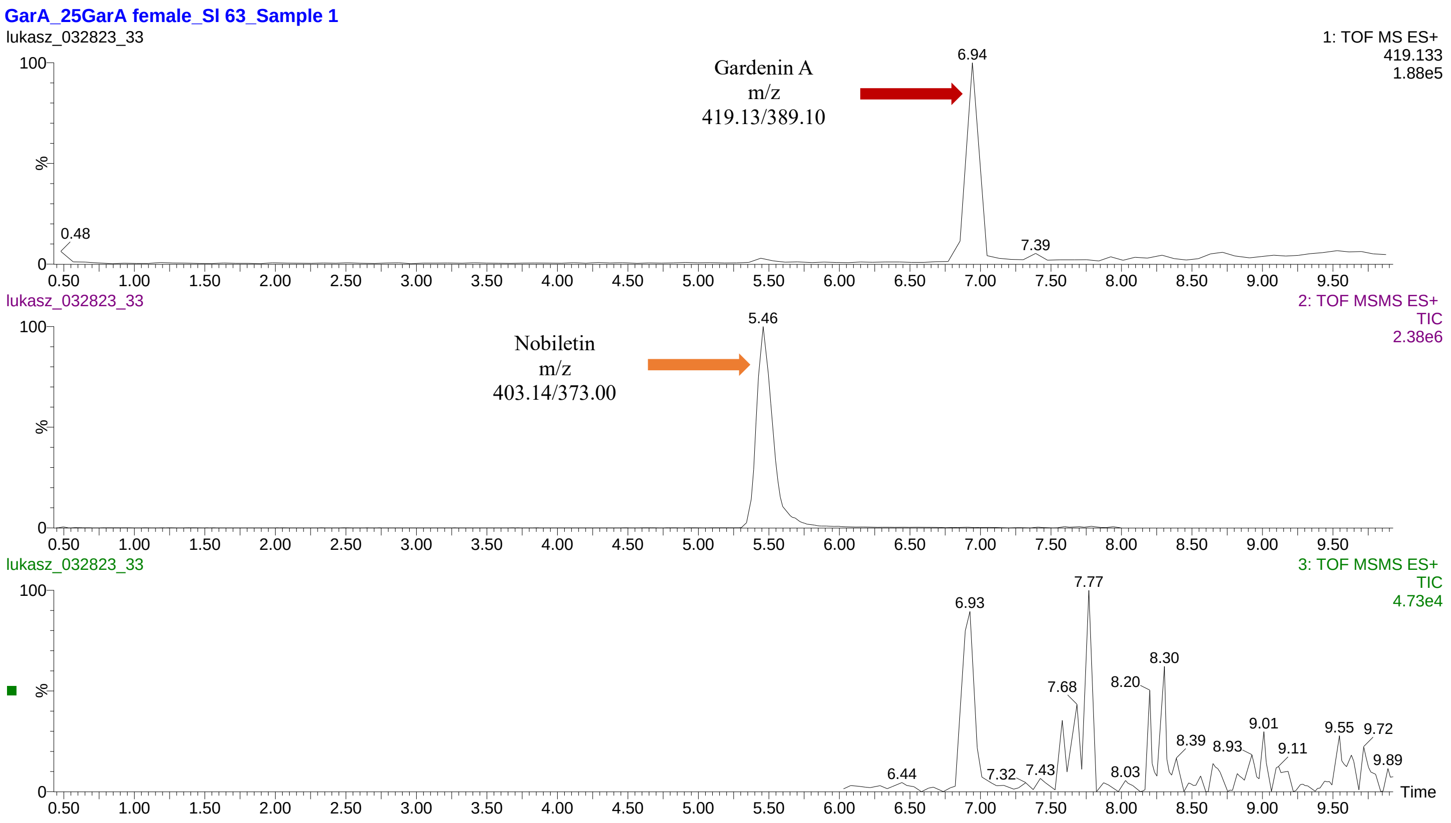
GarA_25GarA female_SI 63_Sample 1
lukasz_032823_33
1: TOF MS ES+
419.133
1.88e5
Gardenin A
m/z
419.13/389.10
6.94
0.48
7.39
lukasz_032823_33
2: TOF MSMS ES+
TIC
2.38e6
Nobiletin
m/z
403.14/373.00
5.46
lukasz_032823_33
3: TOF MSMS ES+
TIC
4.73e4
6.93
7.77
8.30
7.68
8.20
8.39
8.93
9.01
9.11
9.55
9.72
9.89
6.44
7.32
7.43
8.03
100
%
0
0.50 1.00 1.50 2.00 2.50 3.00 3.50 4.00 4.50 5.00 5.50 6.00 6.50 7.00 7.50 8.00 8.50 9.00 9.50
Time

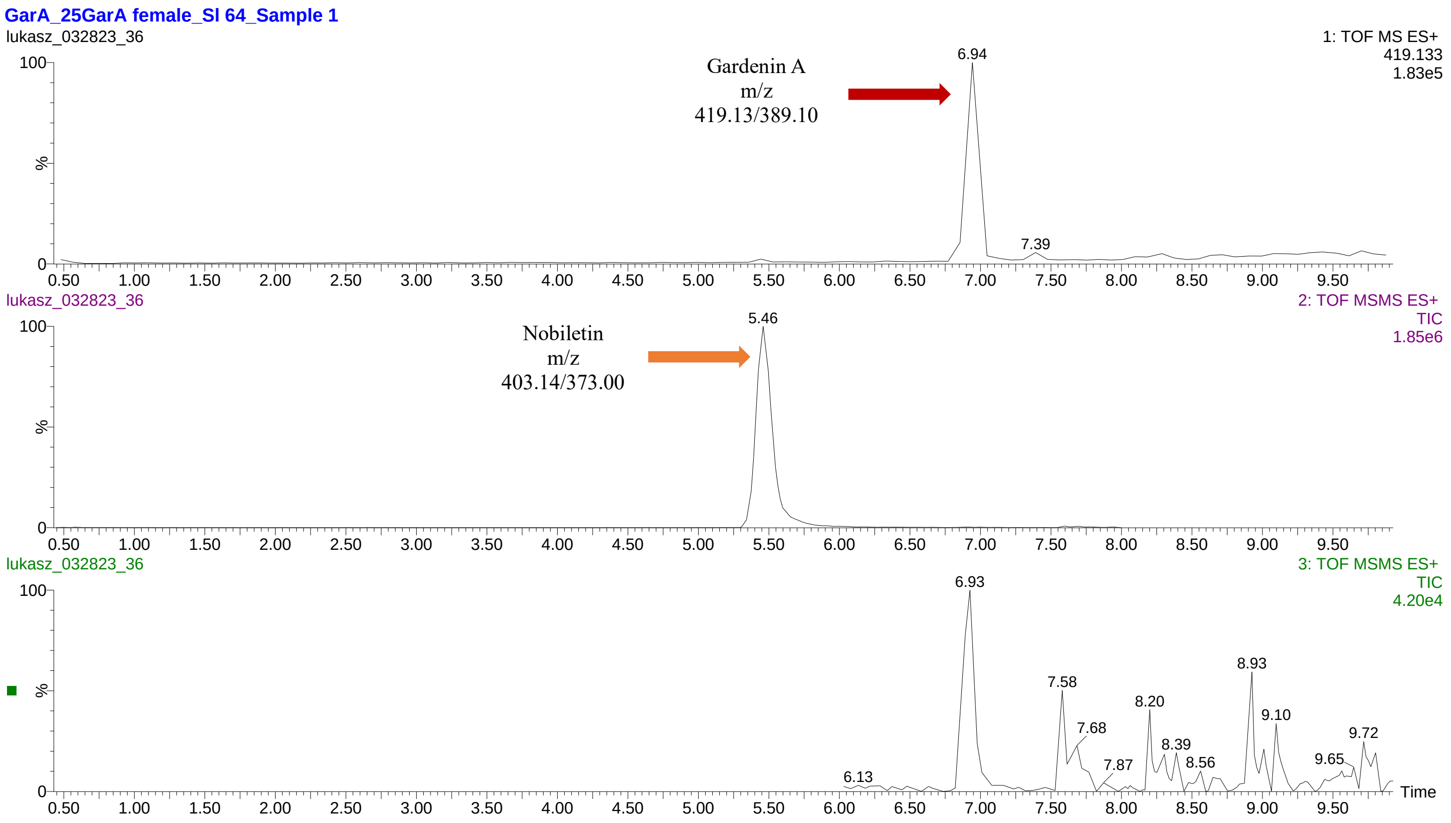
GarA_25GarA female_SI 64_Sample 1
lukasz_032823_36
1: TOF MS ES+
419.133
1.83e5
Gardenin A
m/z
419.13/389.10
6.94
7.39
lukasz_032823_36
2: TOF MSMS ES+
TIC
1.85e6
Nobiletin
m/z
403.14/373.00
5.46
lukasz_032823_36
3: TOF MSMS ES+
TIC
4.20e4
6.93
8.93
7.58
8.20
9.10
7.68
9.72
8.39
7.87
8.56
9.65
6.13
100
%
0
0.50 1.00 1.50 2.00 2.50 3.00 3.50 4.00 4.50 5.00 5.50 6.00 6.50 7.00 7.50 8.00 8.50 9.00 9.50
Time

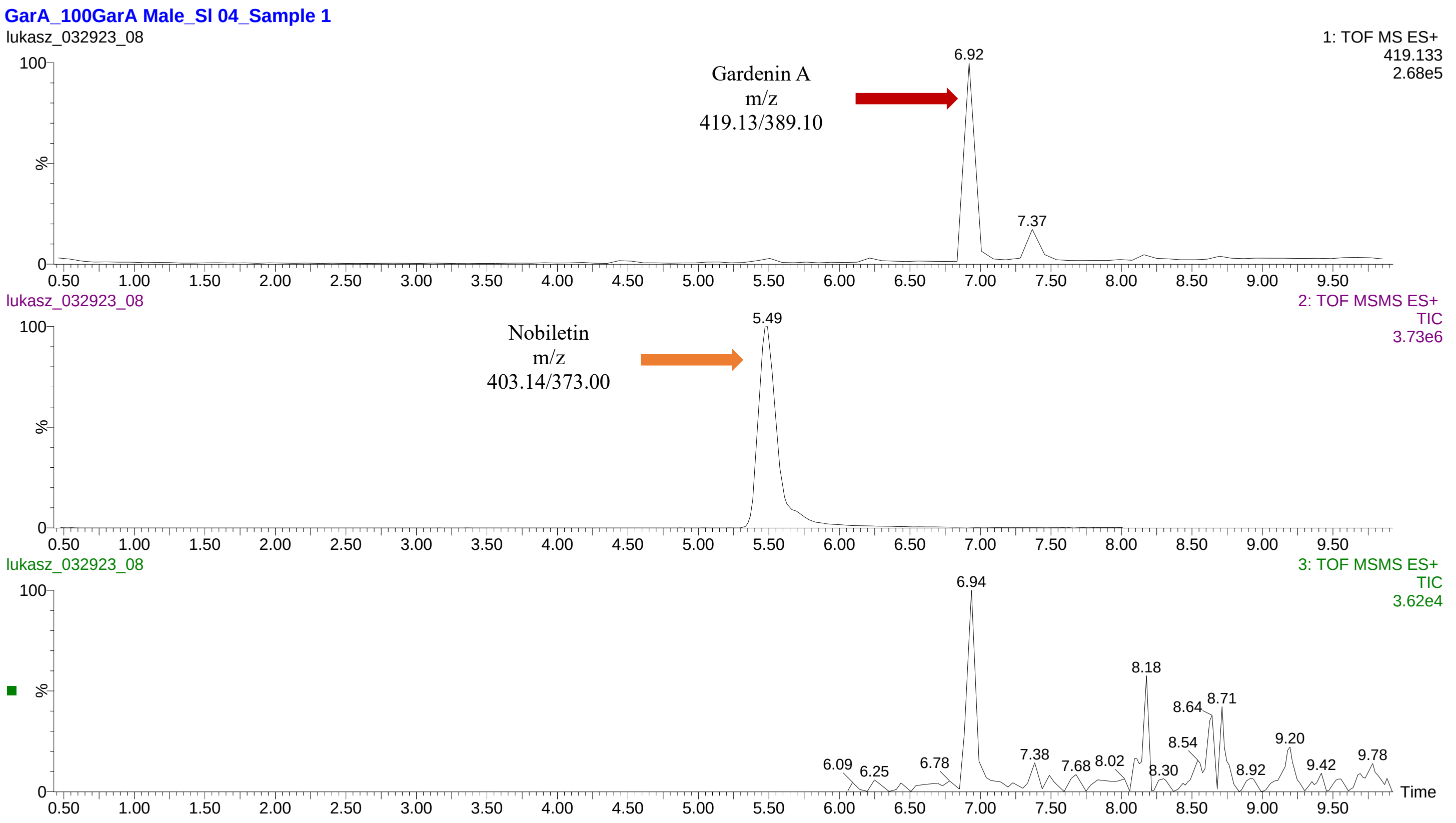
GarA_100GarA Male_SI 04_Sample 1
lukasz_032923_08
1: TOF MS ES+
419.133
2.68e5
Gardenin A
m/z
419.13/389.10
6.92
7.37
lukasz_032923_08
2: TOF MSMS ES+
TIC
3.73e6
Nobiletin
m/z
403.14/373.00
5.49
lukasz_032923_08
3: TOF MSMS ES+
TIC
3.62e4
6.94
8.18
8.64
8.71
8.54
9.20
6.09
6.25
6.78
7.38
7.68
8.02
8.30
8.92
9.42
9.78
%
Time

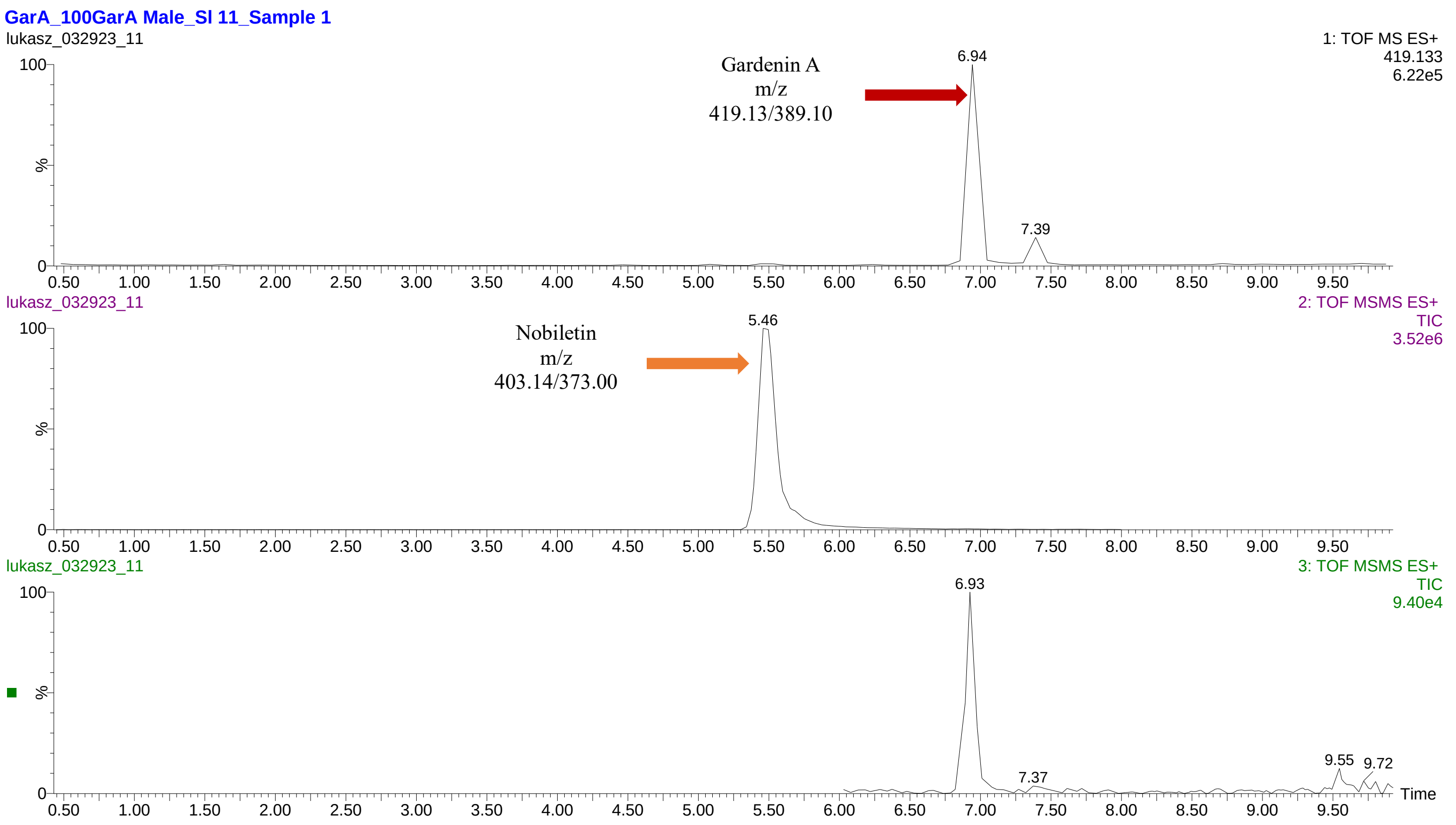
GarA_100GarA Male_SI 11_Sample 1
lukasz_032923_11
1: TOF MS ES+
419.133
6.22e5
Gardenin A
m/z
419.13/389.10
6.94
7.39
lukasz_032923_11
2: TOF MSMS ES+
TIC
3.52e6
Nobiletin
m/z
403.14/373.00
5.46
lukasz_032923_11
3: TOF MSMS ES+
TIC
9.40e4
6.93
7.37
9.55
9.72
%
Time

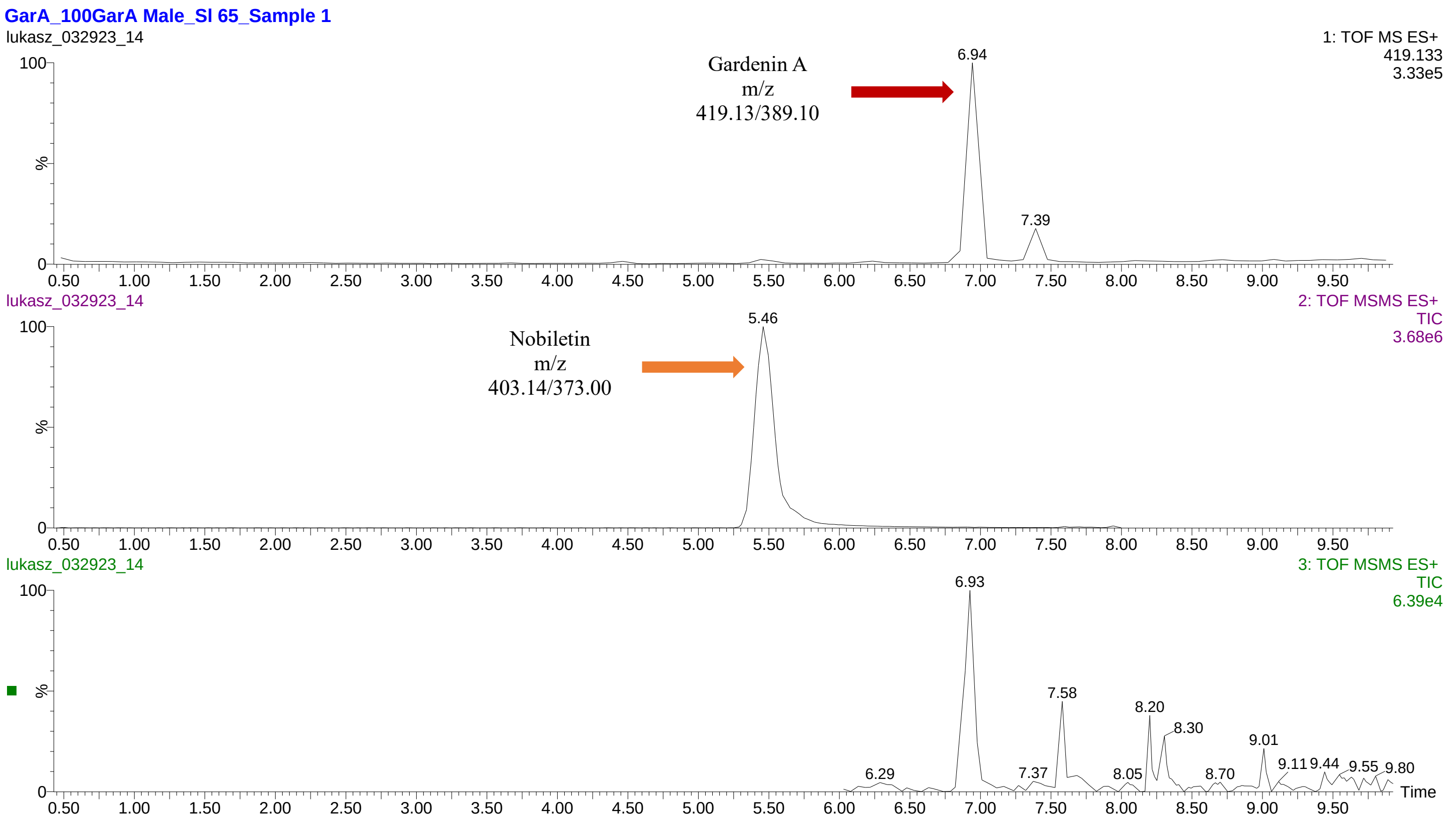
GarA_100GarA Male_SI 65_Sample 1
lukasz_032923_14
1: TOF MS ES+
419.133
3.33e5
Gardenin A
m/z
419.13/389.10
6.94
7.39
2: TOF MS/MS ES+
TIC
3.68e6
Nobiletin
m/z
403.14/373.00
5.46
3: TOF MS/MS ES+
TIC
6.39e4
6.93
7.58
8.20
8.30
9.01
6.29
7.37
8.05
8.70
9.11
9.44
9.55
9.80
%
Time

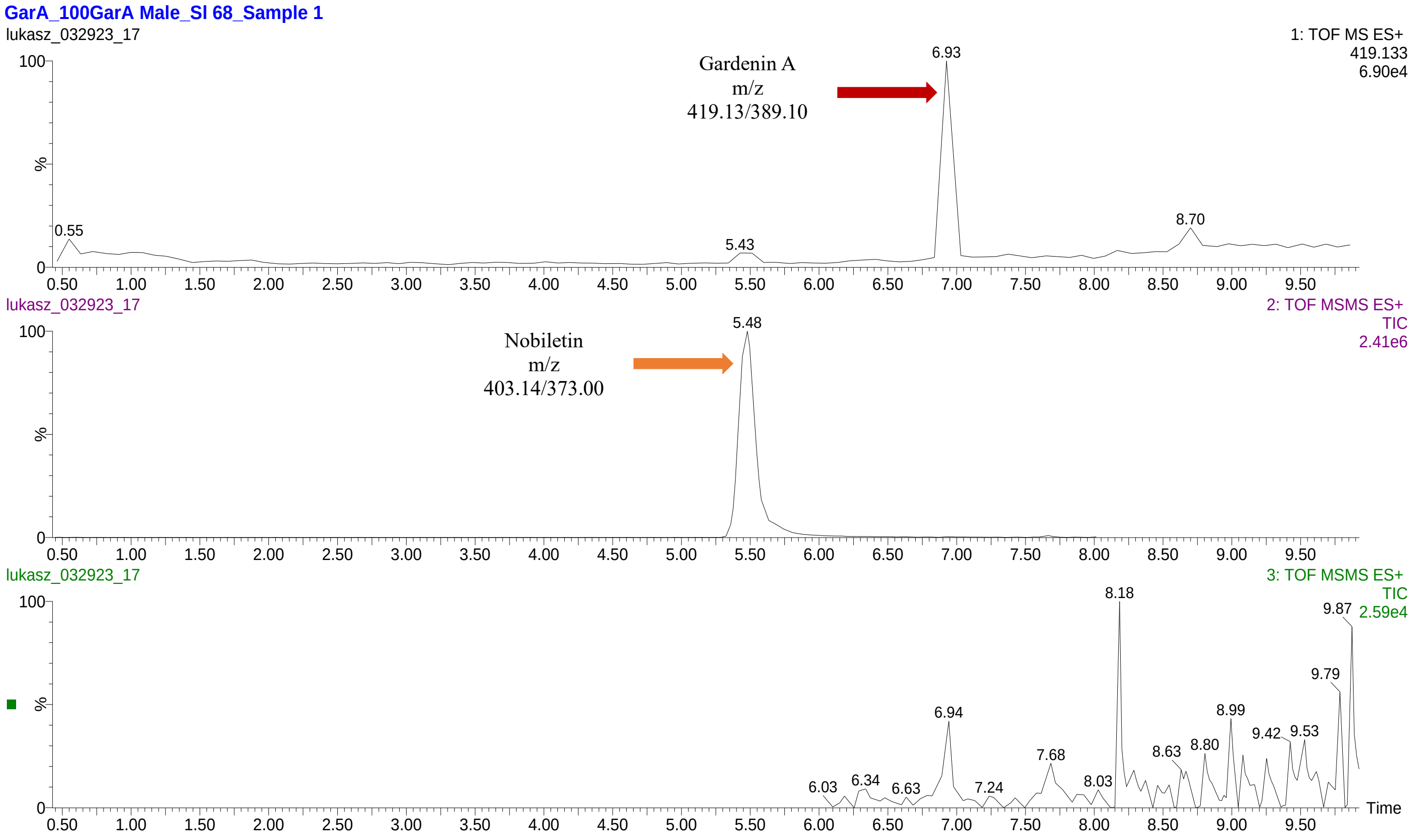
GarA_100GarA Male_SI 68_Sample 1
lukasz_032923_17
1: TOF MS ES+
419.133
6.90e4
Gardenin A
m/z
419.13/389.10
6.93
0.55
5.43
8.70
2: TOF MS/MS ES+
TIC
2.41e6
Nobiletin
m/z
403.14/373.00
5.48
3: TOF MS/MS ES+
TIC
2.59e4
8.18
9.87
9.79
6.94
8.99
9.42
9.53
8.63
8.80
7.68
8.03
6.03
6.34
6.63
7.24
%
Time

**GarA_100GarA Male_SI 69_Sample 1**

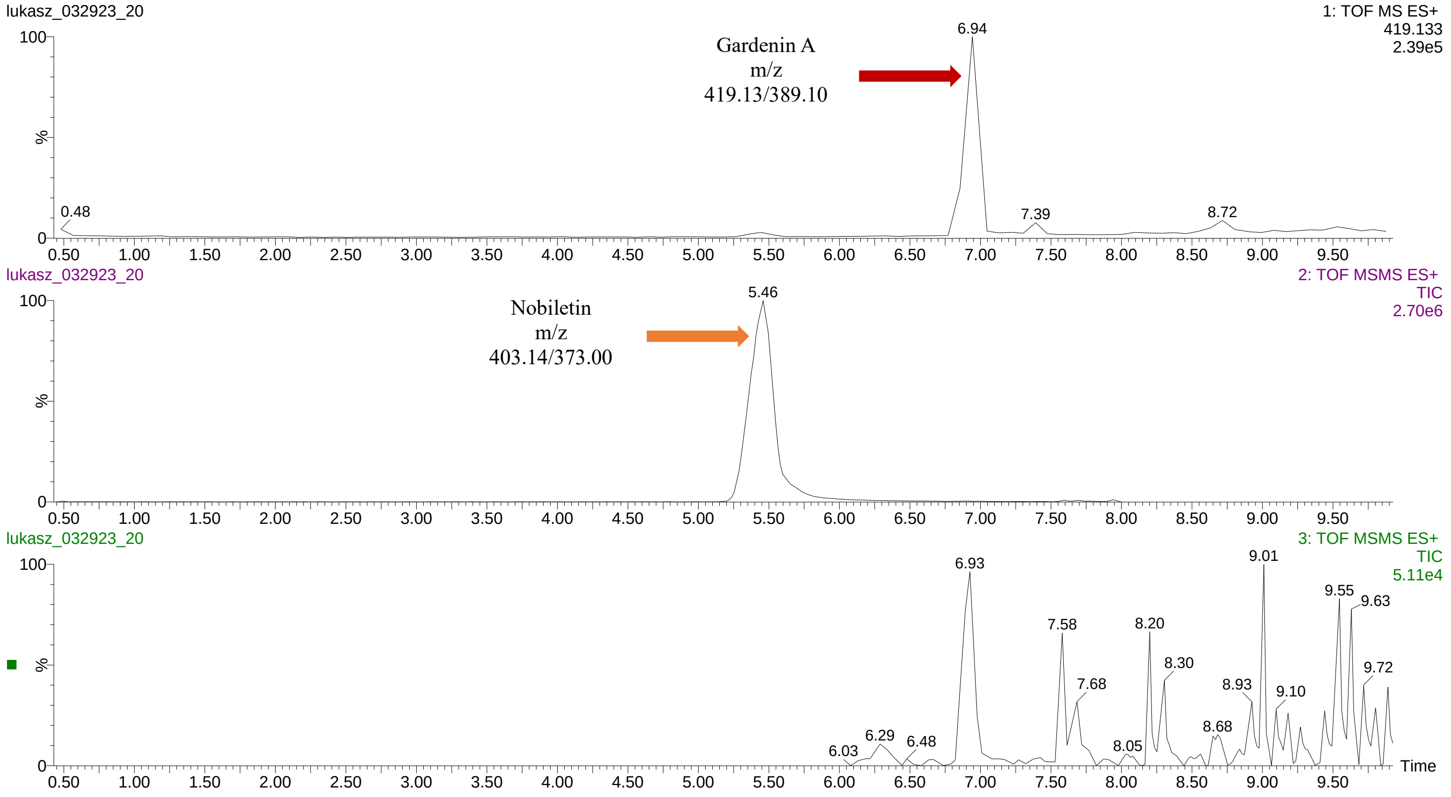


**GarA_100GarA Male_SI 70_Sample 1**

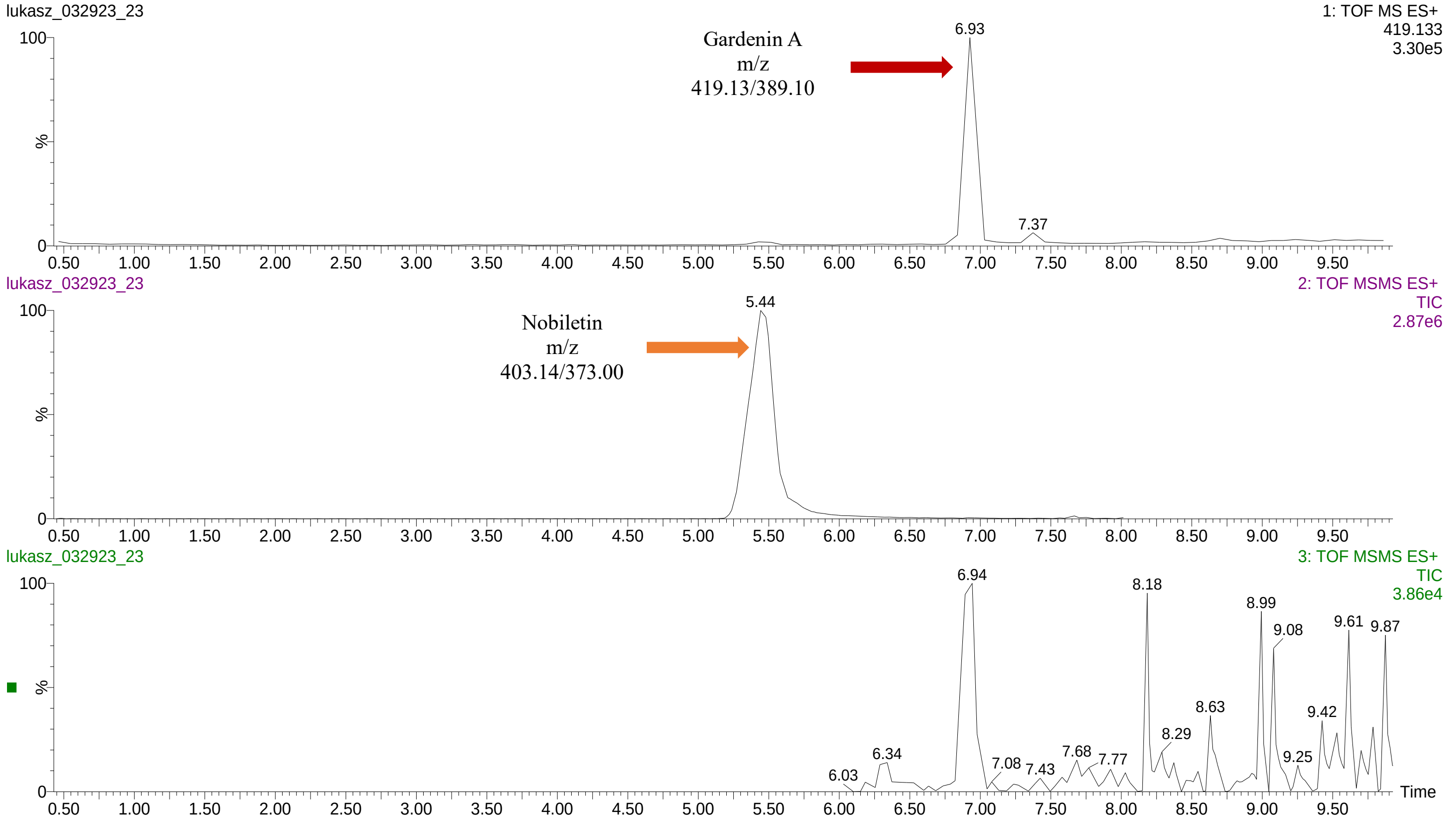

**GarA_100GarA Female_SI 06_Sample 1**

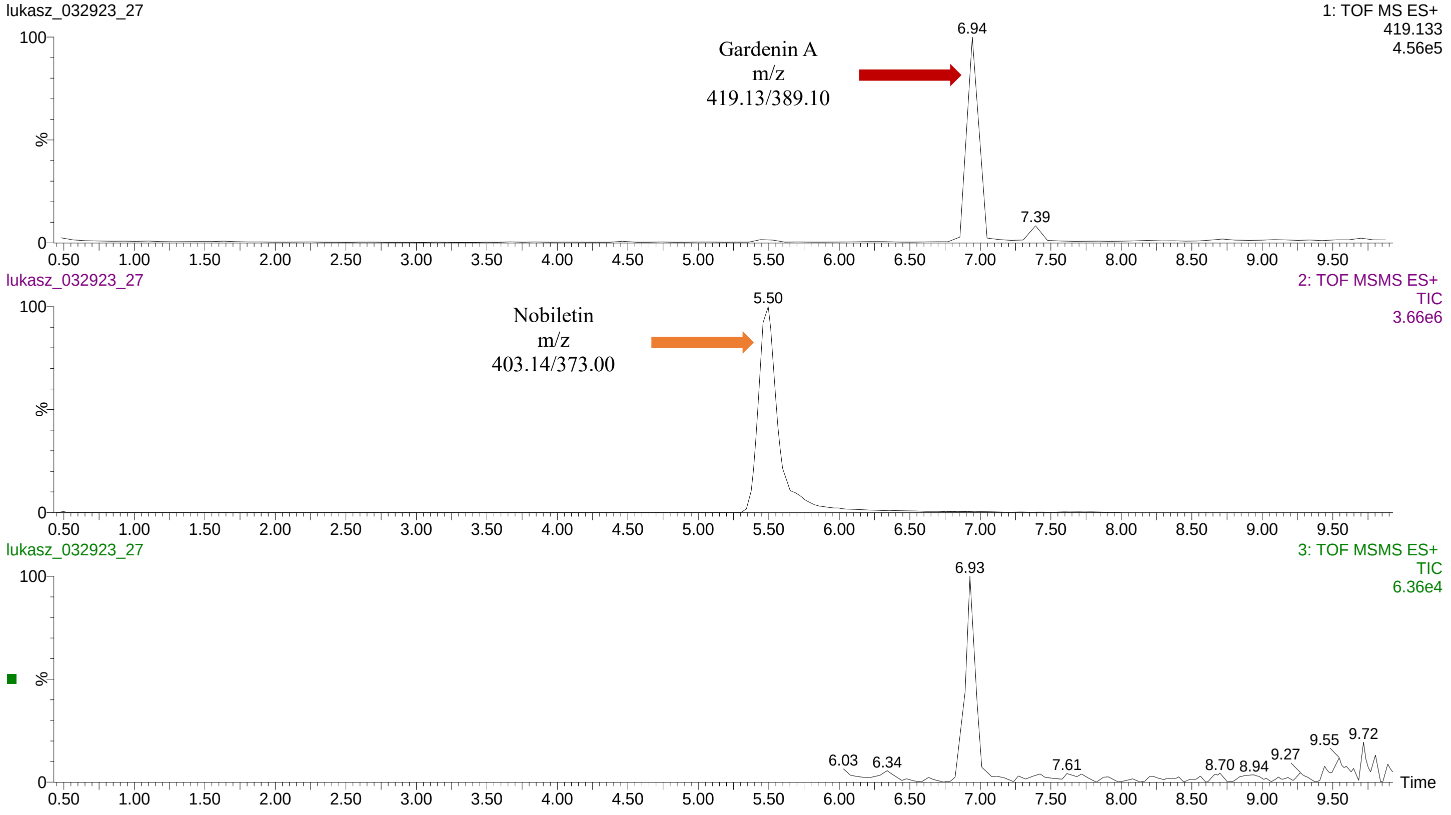


**GarA_100GarA Female_SI 34_Sample 1**

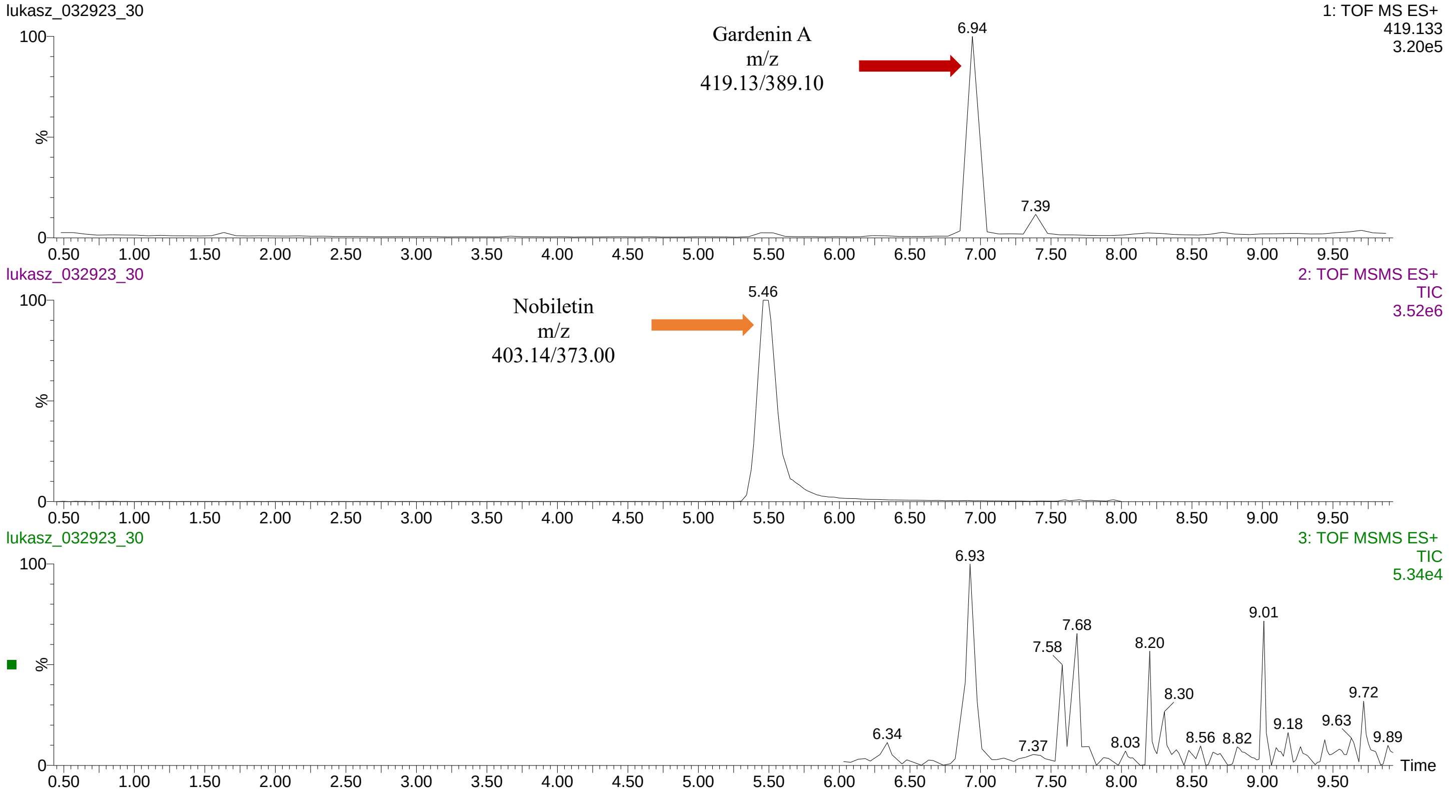

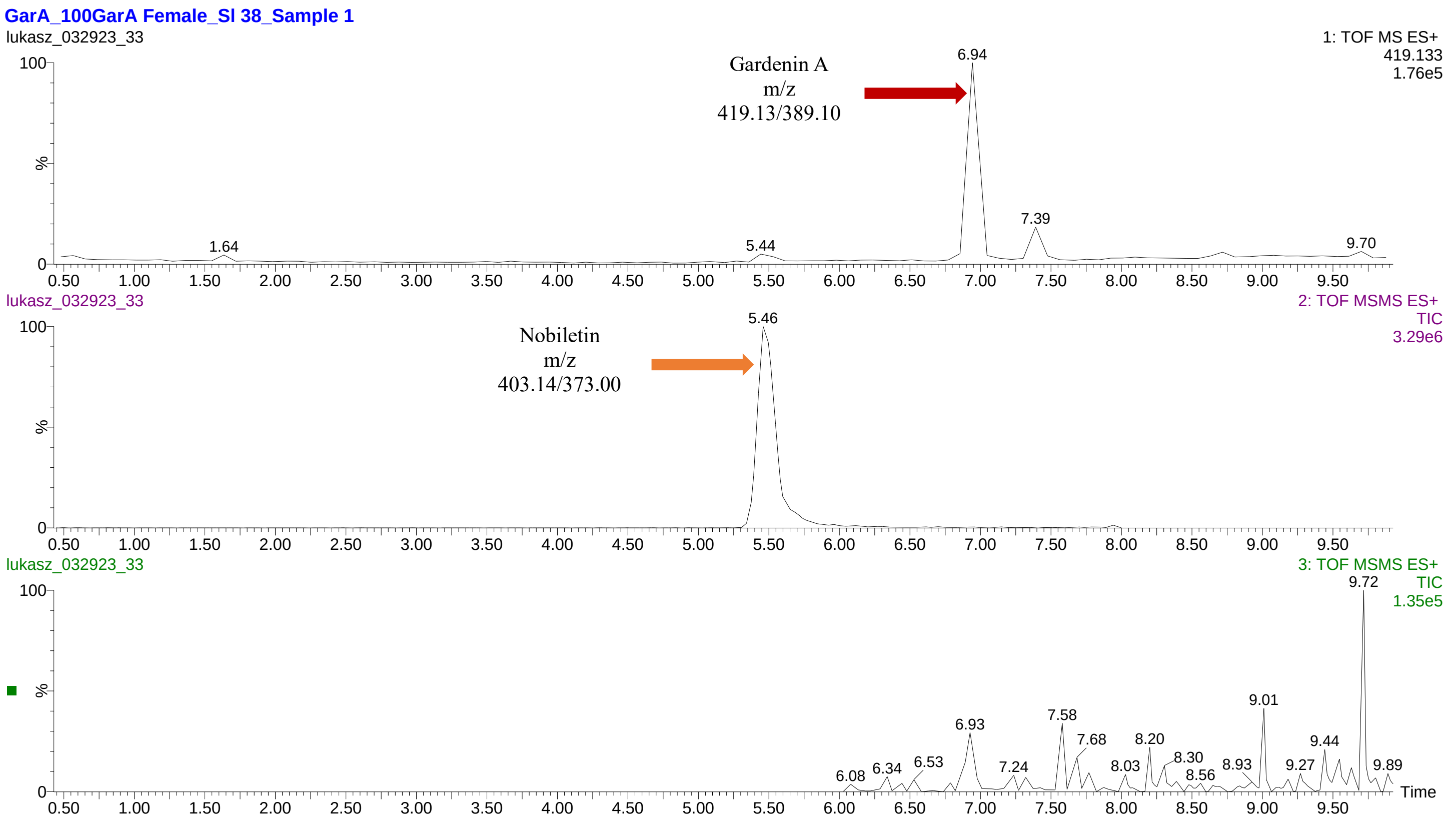
GarA_100GarA Female_SI 38_Sample 1
lukasz_032923_33
1: TOF MS ES+
419.133
1.76e5
Gardenin A
m/z
419.13/389.10
6.94
7.39
1.64
5.44
9.70
lukasz_032923_33
2: TOF MSMS ES+
TIC
3.29e6
Nobiletin
m/z
403.14/373.00
5.46
lukasz_032923_33
3: TOF MSMS ES+
TIC
1.35e5
9.72
9.01
6.93
7.58
7.68
8.20
9.44
8.30
8.56
8.93
9.27
9.89
6.08
6.34
6.53
7.24
8.03
%
Time

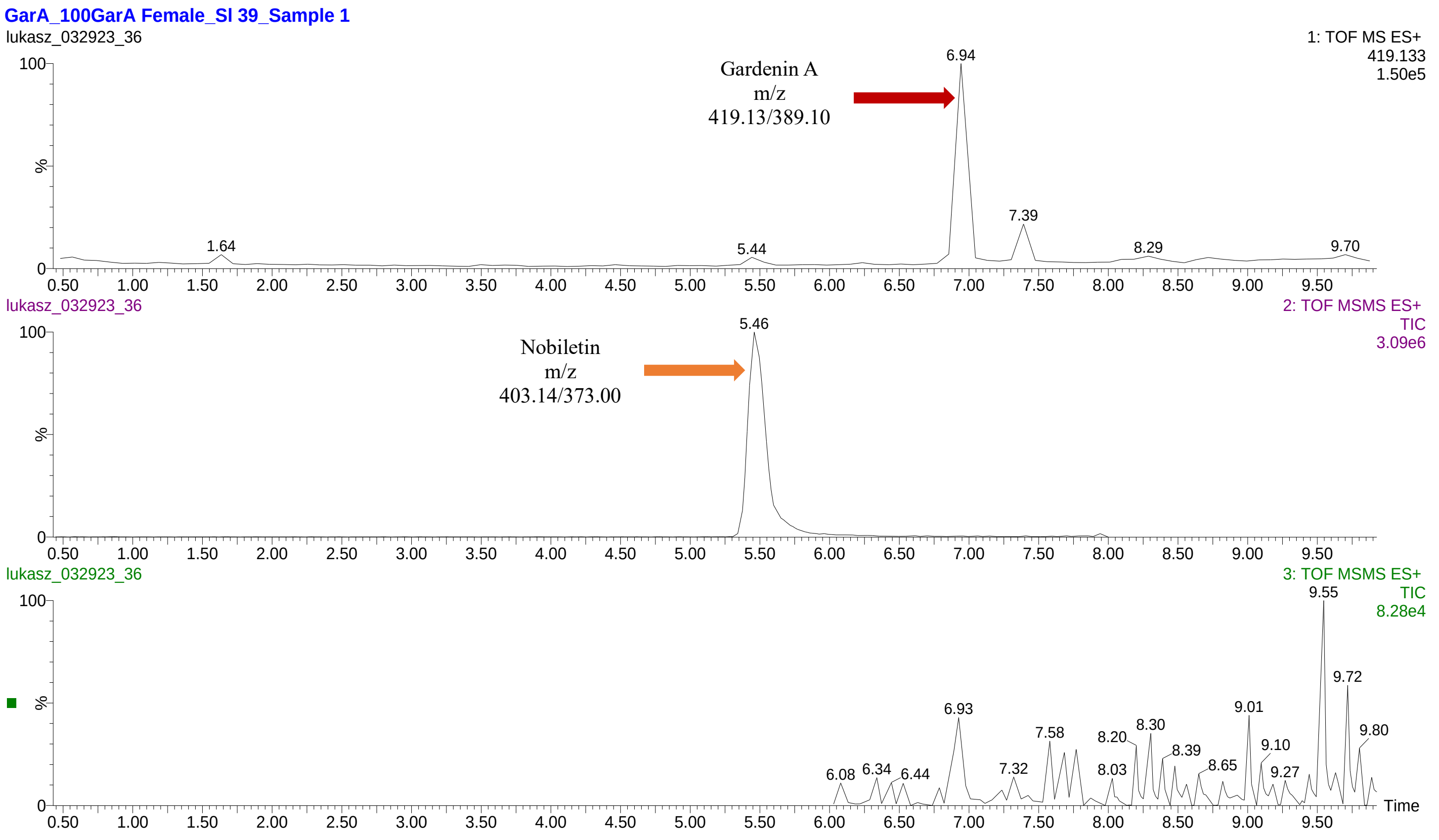
GarA_100GarA Female_SI 39_Sample 1
lukasz_032923_36
1: TOF MS ES+
419.133
1.50e5
Gardenin A
m/z
419.13/389.10
6.94
7.39
1.64
5.44
8.29
9.70
lukasz_032923_36
2: TOF MSMS ES+
TIC
3.09e6
Nobiletin
m/z
403.14/373.00
5.46
lukasz_032923_36
3: TOF MSMS ES+
TIC
8.28e4
9.55
9.72
9.80
9.01
9.10
9.27
6.93
7.58
8.20
8.30
8.39
8.65
7.32
8.03
6.08
6.34
6.44
%
Time

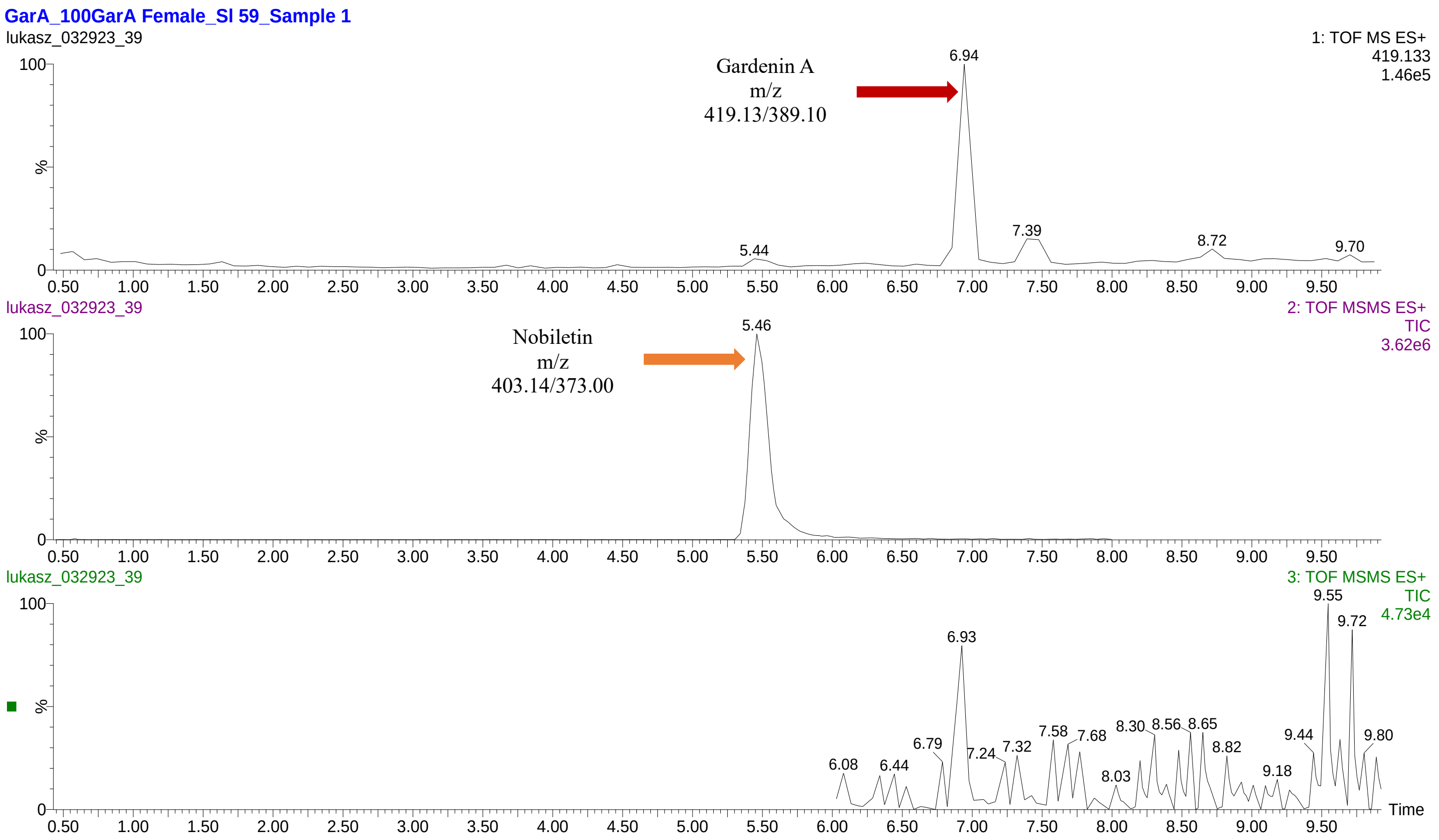
GarA_100GarA Female_SI 59_Sample 1
lukasz_032923_39
1: TOF MS ES+
419.133
1.46e5
Gardenin A
m/z
419.13/389.10
6.94
7.39
5.44
8.72
9.70
lukasz_032923_39
2: TOF MSMS ES+
TIC
3.62e6
Nobiletin
m/z
403.14/373.00
5.46
lukasz_032923_39
3: TOF MSMS ES+
TIC
4.73e4
9.55
9.72
6.93
6.08
6.44
6.79
7.24
7.32
7.58
7.68
8.03
8.30
8.56
8.65
8.82
9.18
9.44
9.80
%
100
0
Time
